# A survey of optimal strategy for signature-based drug repositioning and an application to liver cancer

**DOI:** 10.1101/2021.06.29.450305

**Authors:** Chen Yang, Mengnuo Chen, Siying Wang, Ruolan Qian, Xiaowen Huang, Jun Wang, Zhicheng Liu, Wenxin Qin, Cun Wang, Hualian Hang, Hui Wang

## Abstract

Pharmacologic perturbation projects, such as Connectivity Map (CMap) and Library of Integrated Network-based Cellular Signatures (LINCS), have produced many perturbed expression data, providing enormous opportunities for computational therapeutic discovery. However, currently there is no consensus on which methodologies and parameters are the most optimal to conduct such analysis. Aiming to fill this gap, we developed new benchmarking standards for quantitatively estimating drug retrieval performance. Investigations of potential factors influencing drug retrieval were conducted based on these standards. As a result, we determined an optimal strategy for LINCS data-based therapeutic discovery. With this approach, we further identified new therapeutics for liver cancer of which the current treatment modalities remain imperfect. Both computational and experimental results demonstrated homoharringtonine (HHT) could be a promising anti-liver cancer agent. In summary, our findings will not only impact the future applications of LINCS data but also offer new opportunities for therapeutic intervention for liver cancer.

## 1. Introduction

Despite the major advances in drug research and development (R&D), the cost for *de novo* drug development remains high, ranging from $3 billion to more than $30 billion. Moreover, it usually takes over 10 years to bring a new drug from bench to bedside, reflecting the complex challenges in this area [1]. Within this context, exploring new indications for existing drugs (drug-centric) or identifying effective drugs for certain disease (disease-centric) represent an appealing concept, namely ‘drug repositioning’ (or ‘drug repurposing’), that can greatly shorten the gap between preclinical drug research and clinical applications [2]. Leveraging big data-driven approaches, drug repositioning can be performed computationally, which has the potential to complement traditional therapeutic discovery means and further improve the cost-effectiveness of drug development [3]. The most notably data resources supporting the *in silico*-based therapeutic discovery campaigns would be the Connectivity Map (CMap) [4] and its recent extension called Library of Integrated Network-Based Cellular Signatures (LINCS) [5]. These two projects have generated large-scale drug-induced gene expression profiles on multiple cancer cell lines under different treatment conditions (CMap Build 2: 3 cell lines, 1,309 compounds; LINCS: 77 cell lines, 19,811 compounds), representing a treasure trove for *in silico* therapeutic exploration [6]. As a 1,000-fold scale-up of the original CMap, LINCS contained dramatic increases in both cell line types and perturbations, making it the focus of the present investigation.

The computational drug discovery approach using LINCS (also CMap) data is based upon the basic concept called ‘signature reversion’ [3]. Specifically, to identify candidate agents for treating a certain disease, molecular signature characterizing the states of this disease needs to be generated first. Then, through comparing compound signatures with disease signatures, inverse compound-disease relationships can be identified. Compounds with the ability to reverse disease-specific gene expression pattern are considered as therapeutic candidates (Fig. 1). Several successful applications of this approach include the discovery of rapamycin as combination partner of glucocorticoids in treating lymphoid malignancies [7]; topiramate as a potential agent against inflammatory bowel disease [8]; citalopram as a therapeutic option for patients with metastatic colorectal cancer [9]; niclosamide and pyrvinium pamoate as potential therapeutics for liver cancer [10]. Despite its success, there are still many unsolved problems with this approach. Currently, due to the lack of appropriate benchmarking standards, limited studies have focused on investigating the potential factors that influence this computational strategy, let alone quantitatively evaluating whether these factors affect the accuracy of drug prediction. Therefore, across current signature reversion studies, no consensus regarding the implementation details has been reached. Constructing the rational benchmarking standards and developing the best practice approach can facilitate the development of computational pharmacology area and help to identify more effective therapeutic strategies for refractory diseases.

**Fig. 1.**
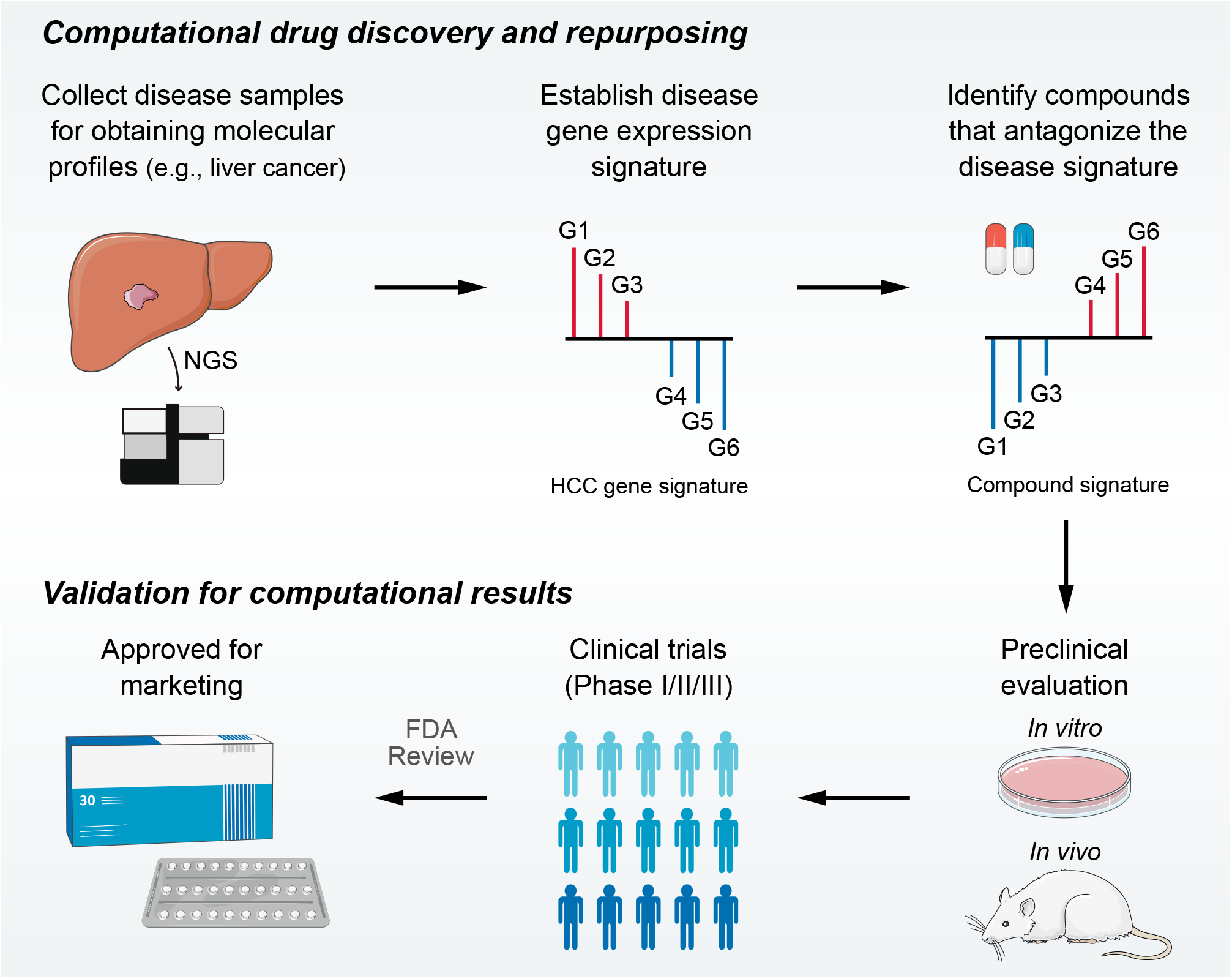
Overview of LINCS data-driven therapeutic discovery. The working principle of ‘signature reversion’-based computational approach. A disease signature representing discordant expression pattern needs first to be identified (G1, G2, and G3 stand for up-regulated genes while G4, G5, and G6 stand for down-regulated genes in disease state). With this signature, pharmacologic perturbation datasets can be queried to find compounds with the ability to reverse disease expression pattern (suppress expression of G1, G2, and G3 and induce expression of G4, G5, and G6). After determining the candidate compounds, experimental and clinical validation are required to translate computational findings to clinical applications.

Herein we mainly focused on the disease of liver cancer. As one of the most lethal malignancies worldwide, liver cancer directly accounts for nearly one million deaths each year (8.2% of new cases of cancer-related death) [11]. Hepatocellular carcinoma (HCC) is the major type of liver cancer, representing approximately 90% of all liver cancer cases [12]. Although many standard of care therapies, including Lenvatinib [13], regorafenib [14], cabozantinib [15], ramucirumab [16], pembrolizumab [17], nivolumab [18] and atezolizumab-bevacizumab [19], have been approved for treating HCC in recent years, most of them can yield only marginal survival benefit. Thus, more effective therapeutics treatments for HCC are highly desired. The objectives of the present study were threefold. The first objective was to develop novel benchmarking standards for evaluating drug retrieval performance. The second one was to determine the best practice approach for LINCS data-based computational drug repositioning. For the last objective, through exploiting the findings from the second objective, we aimed to identify novel drug candidates against liver cancer.

## 2. Results

### 2.1. Summary of influencing factors and compound experiments in LINCS

Many factors may affect the accuracy of signature-based drug retrieval. We have categorized these factors into three main aspects: acquisition of compound signature (reference signature), generation of disease signature (query signature), and selection of disease-compound matching methods (Additional file 2: Figure S1). Although all factors were mentioned and discussed, not all of them were included into the present analyses, considering that some factors have been covered elsewhere and some were challenging to explore due to data and method restrictions. In this study, systematic analyses were carried out to assess the influences of four major factors on signature matching-based drug discovery, including source of cell line, clinical phenotype of query signature, query signature size, and signature matching method. Detailed descriptions and discussions about all included factors were stated in Additional file 1: Supplementary Discussion.

Since only compound-induced expression data was the focus of this study, we first excluded experiments of other perturbagens, including gene knockdown (or knockout) and gene overexpression manipulations. Subsequently, the distribution of compound profiles was visualized based on their perturbation time, perturbation dose, and cell line used. Most of the measurements were made in the treatment durations of 6h (43%) or 24h (56.6%), and under the concentrations of 5μM (21%) or 10μM (63%) (Additional file 2: Figure S2a). The count distribution of all cell lines in LINCS was also presented in Additional file 2: Figure S2b. Although 71 cell lines were included in LINCS project in total, not all of them were extensively profiled, and only nine cell lines contained more than 5,000 profiles, which, however, account for 77.8% of all compound profiles. There were 2,912 compounds shared by these nine cell lines. We further integrating annotation of the most profiled cell lines with treatment duration and concentration information and illustrated the specific profile numbers of each cell line under the conditions of certain time and dose (Additional file 2: Figure S2c). Notably, all the following analyses were performed using LINCS data based on a fixed perturbation condition of 10 μM for 6 hours. Besides, compound profiles of alltop nine cell lines were only utilized when investigating the factor ‘Source of cell line’. In other cases, we focused exclusively on the cell line of HepG2, as in this study, our main point was to uncover novel therapeutics for liver cancer. A systematic summary of included datasets for analyses was presented in Additional file 3: Table S1 and S2.

### 2.2. Cell-type specific gene expression changes induced by compound treatments

Some previous studies utilized compound profiles from cell lines irrelevant to the disease their studied for performing signature-based drug repositioning. To investigate whether this was a reasonable practice, we conducted following analyses based on the LINCS data of the nine most profiled cell lines. First, we visualized the compound profiles in a cosine distance-based two-dimensional t-distributed stochastic neighbor embedding (t-SNE) plot that represented the overall compound perturbation space wherein each dot was equivalent to a unique perturbation and each cell line was color-coded (Fig. 2a). As shown in the figure, most dots with the same color clustered together, indicating that most of compound-induced gene expression changes tended to be cell-type specific. Intriguingly, dots with different colors in the white region seemed to mix together, suggesting that some certain compounds might induce similar gene expression changes across cell lines. To figure out which compounds were likely to cause cell-type specific gene expression changes, and which tended to induce universal changes independent of cell lines, we calculated the pairwise cosine similarities (L1) among the profiles from the same compounds measured in different cell lines (Fig. 2b). The cosine similarity measures range from -1 to 1, where higher values indicate increased similarity. The similarity scores (compound-level, L2) of the 2,912 unique compounds were determined by calculating the median pairwise cosine similarity values (L1) across the most profiled nine cell lines mentioned before (Additional file 3: Table S3). As a result, a high degree of cell-specificity was observed for most compounds, with a median L2 similarity score of 0.078 (Fig. 2c). Furthermore, we retrieved the mechanism of action (MOA) information and mapped them to the compounds to determine the MOA-level similarity scores (L3). L3 similarity scores were calculated based on the median values of L2 similarity scores of compounds within the same MOA. Results suggested that inhibitors targeting core cellular processes (e.g., cell cycle, RNA transcription, protein synthesis) tended to induce similar changes across all cell lines, generally in agreement with previous findings (Fig. 2d and Additional file 3: Table S3) [5, 20]. We then marked the dots representing the compounds of top five MOAs in the t-SNE plot. As might be expected, most of marked dots fell in clusters within the nonspecific region (Fig. 2e).

**Fig. 2.**
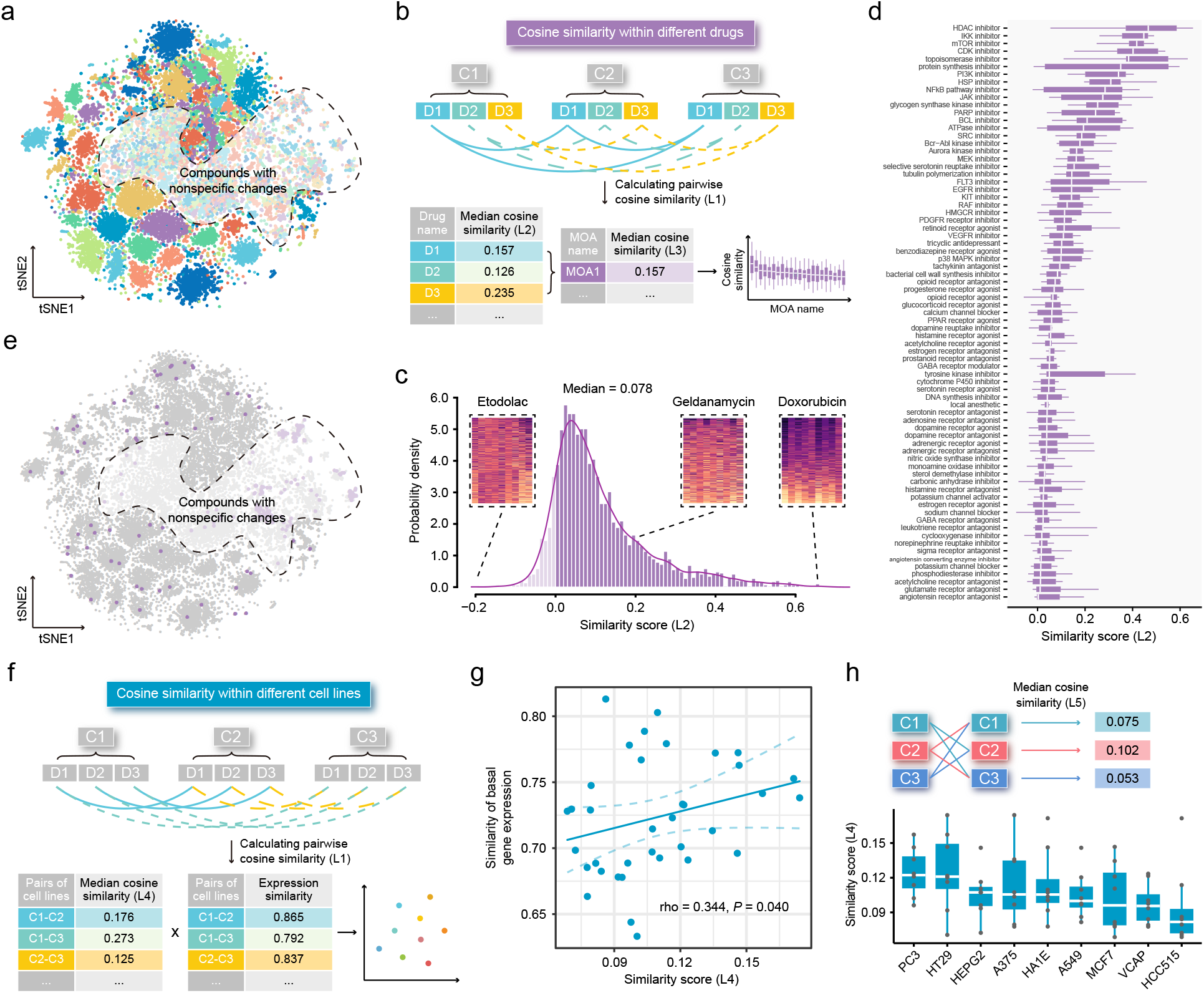
Highly cell-type specific compound-induced expression changes. **a** Two-dimensional t-SNE projection based on cosine distance between compound signatures. Each dot represents a unique perturbation-induced expression profile, and each color represents one type of cell line. **b** Schematic diagram displaying the calculation process of compound-level (L2) and MOA-level (L3) similarity scores. **c** Distribution of compound-level (L2) cosine similarity scores, which ranging from -1 (completely opposite pattern) to 1 (perfectly identical pattern). Three examples are presented (left to right: etodolac, geldanamycin, and doxorubicin). **d** Illustration of MOA-level (L3) similarities. Only MOAs with more than five compounds included are shown in figure. **e** A t-SNE projection showing the distribution of compounds (indicated by purple dots) in top ranked five MOAs (including HDAC inhibitors, IKK inhibitors, mTOR inhibitors, CDK inhibitors, and topoisomerase inhibitors). **f** Schematic diagram displaying the calculation process of cell line pair-level (L4) similarity scores. (**g**) Correlation between basal expression similarities and perturbed expression similarities (L4) of 36 cell line pairs (nine cell lines in total). Statistical significance and correlation coefficient were determined by ranked-based Spearman correlation. **h** Schematic view of the calculation of cell line-level (L5) similarity scores (upper) and the presentation of L5 similarity scores of nine cell lines in the boxplot (lower).

Apart from investigating similarity of perturbed expression profiles at compound level, we sought to further investigate the cell line pair/cell line-level similarity. Nine cell lines contributed a total of 36 unique cell line pairs. The cell line pair-level perturbed expression similarities (L4) were determined through calculating the median value of similarity scores of all compound pairs between two cell lines, and the corresponding basal expression similarities were computed using Spearman ranked correlation on expression data from CCLE project (Fig. 2f). Based on these 36 cell line pairs, we found that there was a significant, albeit not very remarkable, association between the perturbed expression similarities (cell line pair-level, L4) and basal expression similarities (rho = 0.344; *P* = 0.040), suggesting that cell lines with similar molecular features were more likely to have consistent gene expression changes upon perturbation (Fig. 2g). Similarities within the nine cell lines were also explored (cell line-level, L5). Among the nine cell lines we tested, PC3 cell line showed the highest L5 similarity score (median value = 0.122) (Fig. 2h). Notably, the cosine similarity of 0.122 still denoted a weak relationship, which further supported the conclusion that compound-induced gene expression changes were highly cell line specific.

Among the nine most profiled cell lines, HepG2 was the only one derived from liver. To investigate whether HepG2 was an appropriate cell line model for computational therapeutics discovery for liver cancer or other liver-associated diseases, we calculated the expression correlation between HepG2 and other cell lines (921 CCLE cell lines) or tissues (17,382 normal tissues from GTEx and 9,701 tumor tissues from TCGA PanCancer). Compared to other tissue-derived cancer cell lines or normal/tumor tissues, HepG2 exhibited a significantly higher expression correlation with liver cancer cell lines (median correlation coefficient = 0.729), normal liver tissues (median correlation coefficient = 0.616), and liver cancer tissues (median correlation coefficient = 0.631) (Additional file 2: Figure S3). Coupled with above finding of the significant relationship between perturbed expression similarities and basal expression similarities, we supposed that the use of LINCS-derived HepG2 data was preferable to be limited within liver diseases.

### 2.3. Developing new benchmarking standards

Owing to the lack of benchmarking standards, accurate assessment of retrieval performance of signature matching methods remains challenging. Inspired by previous findings [10, 21], we proposed two novel benchmarking standards, namely area under curve (AUC)-based standard and Kolmogorov-Smirnov (KS) statistic-based standard. They were built upon different notions and thus independent of each other, which helped to avoid potential bias introduced by single standard. The corresponding benchmarking datasets were developed mainly based on preclinical/clinical data of liver cancer. Detailed processes regarding data collection and metrics calculation have been described in Methods and visualized in Additional file 2: Figure S4a.

The development of AUC-based standard was based on the finding that there existed correlation between the reversal potency and treatment efficacy, which has been reported by several previous studies [10b, 21a]. In order to further validate whether this correlation remained significant in other conditions, we retrieved drug response data from CTRP dataset in which area under the dose-response curve (AUDRC) values were used as measurements of drug sensitivity, and utilized two different HCC signatures as query signatures to obtain KS-based similarity scores [10]. A total of 109 compounds shared by two datasets were selected to conduct correlation analyses. As a result, statistically significant correlation could still be observed between similarity scores and AUDRC values in these scenarios, further proving the reliability of this standard (Additional file 2: Figure S4b). A benchmark dataset was then generated, composing of 117 unique compounds with both LINCS and drug efficacy (IC_50_) data available, which was taken as a basis for the application of AUC-based standard (Additional file 2: Table S4). The resultant AUC from this standard were termed as drug retrieval-associated AUC (DR-AUC). Higher DR-AUC value indicated better performance.

As for KS statistic-based standard, we assumed that agents under evaluation in clinical trial for HCC treatment, namely HCC agents, might possess an increased reversal capacity [10a]. In other words, HCC agents were more likely to cause negative enrichment in KS test. To verify this hypothesis, we compiled a set of 27 potential HCC agents which were both included in LINCS and under clinical trials for liver cancer treatment. Besides, similarity scores of all compounds tested in HepG2 were also calculated, which were then used as ranked list for KS test. The results of KS test demonstrated the above assumption that the HCC agent set was negatively enriched significantly (Additional file 2: Figure S4c). Accordingly, KS statistic-based standard could be implemented based on this 27-agent set and the resultant enrichment scores (ES) here were termed as drug retrieval-associated ES (DR-ES) (Additional file 3: Table S5). Of note, in contrast to DR-AUC, lower DR-ES values denoted better performance.

### 2.4. Benchmarking signature matching methods for drug retrieval

These two independent benchmarking standards enabled us to quantitatively assess the retrieval performance of different signatures matching methods and thus aided in the identification of optimal methodology for computational drug repositioning. In total, six available methods including eXtreme Sum (XSum) [21b], eXtreme Cosine (XCos) [21b, 22], eXtreme Pearson (XCor) [23], eXtreme Spearman (XSpe) [23], Kolmogorov-Smirnov (KS) test [4], and the Reverse Gene Expression Score (RGES) [10b] were included for performance comparison. To minimize technical bias introduced by different query signatures, four HCC signatures with different sizes generated from distinct datasets were utilized for benchmarking (Additional file 3: Table S6). Of these, Sig_gastro_ [10a] and Sig_NC_ [10b] were directly obtained from previously publications, while Sig_LIRI_ and Sig_GSE54236_ were generated using RNAseq data from LIRI cohort and microarray data from GSE54236, respectively. A brief summary of above essential components involved in the evaluation process was illustrated in Fig. 3a.

**Fig. 3.**
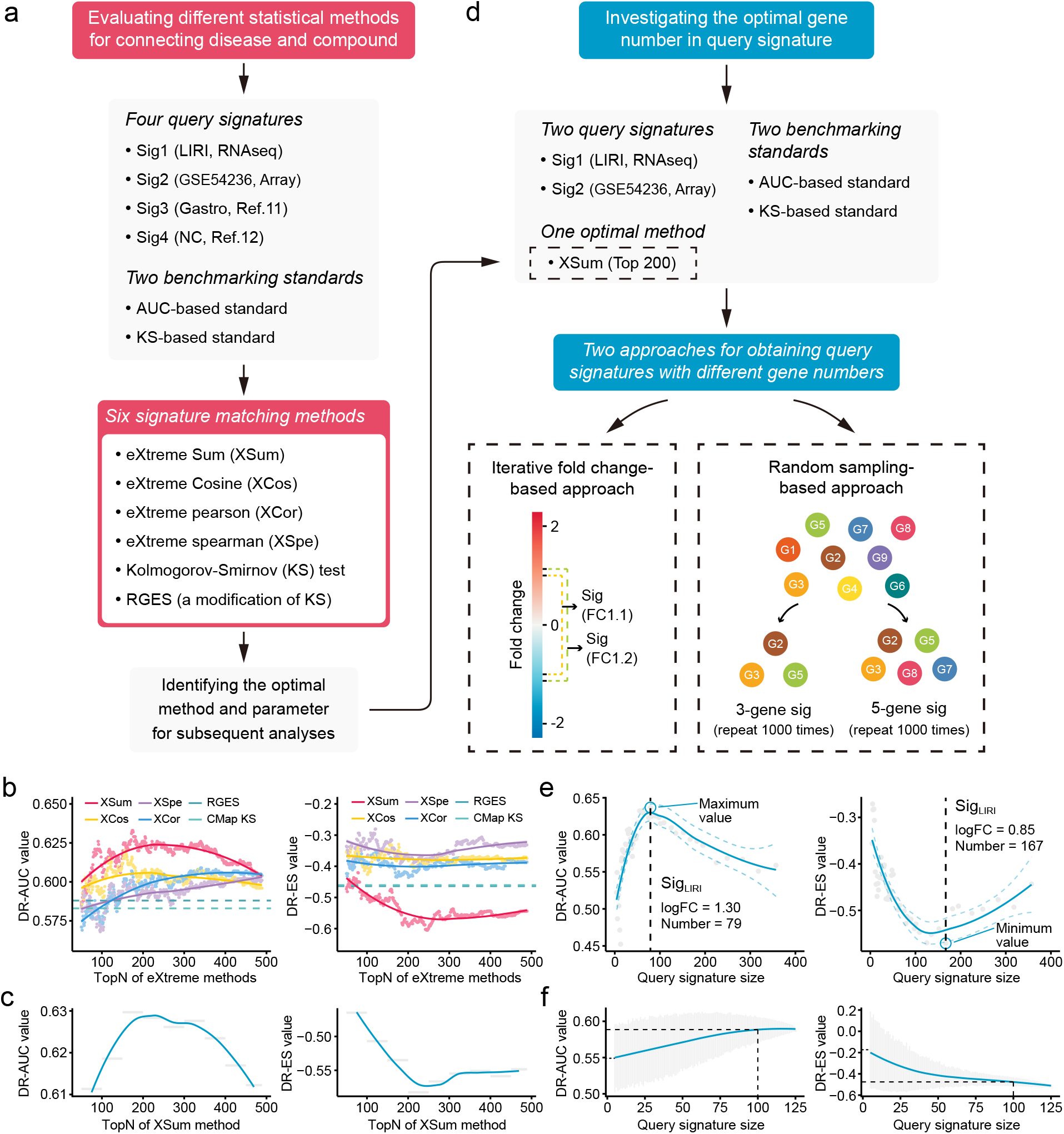
Benchmarking different methodologies and parameters to determine the optimal computational strategy. **a** Diagram summarizing the workflow and the important components involved in the evaluation process of drug retrieval performance of six different signature matching methods. **b** Retrieval performance of six matching methods evaluated by AUC-based benchmarking standard (left) and KS statistic-based benchmarking standard (right). **c** Visualization of AUC-based (left) and KS statistic-based (right) performance measurements of XSum method on standardized data for discerning the optimal operating parameter. **d** Diagram summarizing the workflow and the important components associated with the investigation process of the optimal query signature size. **e** Relationship between query signature size determined by iterative fold change-based approach and retrieval performance evaluated by AUC-based standard (left) and KS statistic-based standard (right). **f** Relationship between query signature size determined by random sampling-based approach and retrieval performance evaluated by AUC-based standard (left) and KS statistic-based standard (right). LOESS polynomial regression analysis was performed for curve fitting. Query signature for above analyses was generated based on LIRI cohort, and evaluation results using other query signatures were presented in Supplementary Figure 5.

Considering that the performance of eXtreme methods (including XSum, XCos, XCor, and XSpe) may be affected by the number of top genes (topN), we thus calculated the DR-AUC or DR-ES values of each eXtreme methods iteratively, using topN ranging from 50 to 489. In the condition of using Sig_LIRI_ as query signature, both benchmarking standards demonstrated that XSum outperformed other five methods across almost all candidate topNs (Fig. 3b). Concordantly, when using other three query signatures, XSum also achieved better performance compared with other methods, except in the case of using Sig_Gastro_ as query signature and AUC-based standard for benchmarking, where RGES showed a similar performance with XSum (Additional file 2: Figure S5a-c). Generally, XSum exhibited a consistently satisfactory performance, independently of the query signature and benchmarking standard, and thus could be used as the optimal methodology for drug retrieval (Additional file 3: Table S7 and S8). In addition, our analyses also demonstrated that the recently developed RGES (a modification of the KS method) appeared superior to the KS method in most cases and might serve as an alternative approach for KS-based connectivity mapping [10b].

Since we have proved that XSum could be used as the optimal matching method, we next sought to find the most appropriate topN value for applying XSum method to achieve the best retrieval performance. Directly selecting the exact topN value where corresponding DR-AUC/DR-ES reached their maximum/minimum might cause bias and could be prone to over fitting. Given the continuous trait of candidate topNs, we chose to divide them into several smaller groups, typically 50 topN values in each group. The DR-AUC/DR-ES of a certain group was defined as the mean DR-AUC/DR-ES values within this group, which could decrease potential influences brought about by the outliers. With this normalization approach, relatively consistent results across all conditions could be obtained. The optimal topN group which yielded best performance were either ‘top150-200’ or ‘top200-250’ groups (Additional file 3: Table S9). Besides, we also observed a biphasic pattern of fitting curves, with the inflection points appearing where topNs were around 200 (Fig. 3c and Additional file 2: Figure S5d-f). Based on these results, we supposed that topN of 200 could serve as a rough guide.

### 2.5. Determining the optimal query signature size

Above analyses determined XSum as the optimal method and tuned the parameter of topN to 200. Next, we intended to further discern the optimal gene number in query signatures. Considering that Sig_gastro_ (n=44) and Sig_NC_ (n=73) had fixed and relatively small signature sizes, we only utilized signatures generated from LIRI and GSE54236 cohorts for subsequent investigation. We adopted two complementary approaches: (i) iterative fold change-based and (ii) random sampling-based approaches, to obtain query signatures with different sizes (Fig. 3d). The iterative fold change-based approach could create a number of signatures with discontinuous sizes through setting iterative threshold values of fold changes. The exact sizes of optimal signatures identified by this approach varied substantially (including 55, 79, 140, and 167). Despite this, similar trends of biphasic pattern with inflection points at around 100 under different conditions could still be observed (Fig. 3e and Additional file 2: Figure S6a). The approach based on random sampling was adopted as a complement. The results showed that as the signature sizes increased, the DR-AUC/DR-ES values also increased/decreased and eventually converged when the signature size was more than 100 (Fig. 3f and Additional file 2: Figure S6b). Based on these observations, we considered that the signature size of 100 could be selected as a good compromise. This conclusion remained valid in the conditions when other topN values were applied, such as 100 and 400 (results not shown).

### 2.6. Investigating the optimal clinical phenotype underlying query signature

Many previous studies chose to compare normal versus diseased states to define disease signatures. However, signatures derived from other clinical phenotypes, such as prognosis and metastasis, could also be used to query LINCS. Aiming to figure out whether this factor could also affect the performance of drug retrieval, we designed a forward and a backward strategy respectively (Additional file 2: Figure S7a). The application of forward strategy was based on two types of signatures, general signatures (representing discordant expression pattern between normal and tumor tissues) and prognostic signatures (associated with survival outcomes). Through adopting above random sampling-based approach, we compared the above two signature phenotypes across varied signature sizes. Unfortunately, the results under different datasets and benchmarking standards were highly inconsistent. This strategy, consequently, failed to provide a definitive conclusion (Additional file 2: Figure S8 a and b).

Opposed to the forward strategy, backward strategy started from creating a collection of 10,000 random signatures, followed by identification of the optimal signature for clinical implication evaluation. Specifically, the optimal random signature was determined according to both benchmarking standards, and exploring the clinical values of this signature might, to some extent, reveal some necessary features possessed by a ‘good’ query signature (Additional file 2: Figure S7b). A comprehensive clinical evaluation on the optimal signature was carried out based on five RNAseq and five microarray clinical cohorts from three perspectives. First, the ability of this signature to distinguish tumors from non-tumors was investigated. Briefly, we extracted the first principal components (PC1) of this signature to represent its overall expression pattern. AUC was used here as a reflection of the classification capability. The results showed that more than 0.90 of AUC can be obtained in seven out of eight cohorts (87.5%), indicating that the ability to discern the difference between diseased and normal states might be an indispensable property for achieving good retrieval performance (Additional file 2: Figure S7c). Next, we intended to find out whether the optimal signature should be a prognostic indicator. Cox regression analyses were conducted to investigate the association between the signature expression (PC1) and clinical outcome. As a result, significant prognostic implications (Cox *P* < 0.05) of the optimal signature could be observed in six out of eight cohorts (75%), suggesting that prognostic significance could also be a necessary characteristic (Additional file 2: Figure S7d). At last, the association between signature expression and other clinical features was explored. Considering that CHCC and LIHC cohorts held the most abundant clinical information, corresponding analyses were thus conducted on these two cohorts. The results showed that there was a significant correlation between the optimal signature and multiple clinical features, including BCLC stage (*P* = 0.011), tumor thrombus (*P* = 0.001), AFP level (*P* = 0.022), TNM stage (*P* < 0.001), and histologic grade (*P* < 0.001) (Additional file 2: Figure S7e). According to the above results, we concluded that quality query signature should possess the ability to comprehensively recapitulate the clinical features of corresponding disease, rather than only reflect the disease characteristic from single perspective.

### 2.7. Linking above findings to the discovery of new anti-liver cancer therapeutics

The knowledge acquired from above analyses was then applied to establish a signature characterizing liver cancer initiation and development, so as to identify computationally prioritized compounds with potential therapeutic as well as chemopreventive effects against liver cancer. The generation of this evolution-associated signature was based on the notion that the initiation and progression of liver cancer was a stepwise process with gradually acquired advantageous biological capabilities (Fig. 4a). Therefore, conceptually, antagonizing genes that were most related to these stages could be a potential therapeutic strategy. Through implementing Random Forests algorithm on GSE89377 cohort, preliminary screening was performed to include stage-associated genes, where genes with greater predictive power were selected for further analysis. This screening yielded a total of 6017 stage-associated genes (23.9%), of which 309 were landmark genes (Fig. 4b). Next, we conducted weighted gene co-expression network analysis (WGCNA) to obtain co-expressed modules with diverse expression patterns (Additional file 2: Figure S9a). Seven gene modules were discerned by WGCNA analysis (Additional file 2: Figure S9 b and c), and two of them, which we termed the ‘ascending’ module (n = 1,738) and the ‘descending’ module (n = 350) for their greatest relevance to stages and patterns of linear evolution from normal to cancer, were retained for further analyses (Fig. 4 c and d). Biological processes associated with genes in these two modules were investigated. We found that the ‘ascending’ module was closely associated with proliferation (Fig. 4c), while the ‘descending’ module was enriched in several different types of processes (Fig. 4d). There were 159 genes in common between these two modules and landmarks. Based on the aforementioned recommend about the optimal query signature size, we sought to further reduce the size of 159 to 100 genes. This procedure was carried out using HCC occurrence-related clinical and molecular data in GSE15654 cohort. In brief, ten thousand random signatures, each containing 100 genes, were generated based on the 159-gene panel. The one which had the most significant association with HCC occurrence was considered as the optimal query signature (Additional file 2: Figure S10 a and b). This analysis yielded a signature comprised 82 ascending genes and 18 descending genes, with cox *P* value of 0.009, which was then termed as Sig_evo_ (Additional file 3: Table S10). The linear evolution pattern of Sig_evo_ remained present in training (Additional file 2: Figure S10c) as well as an internal validation cohort (Additional file 2: Figure S10d).

**Fig. 4.**
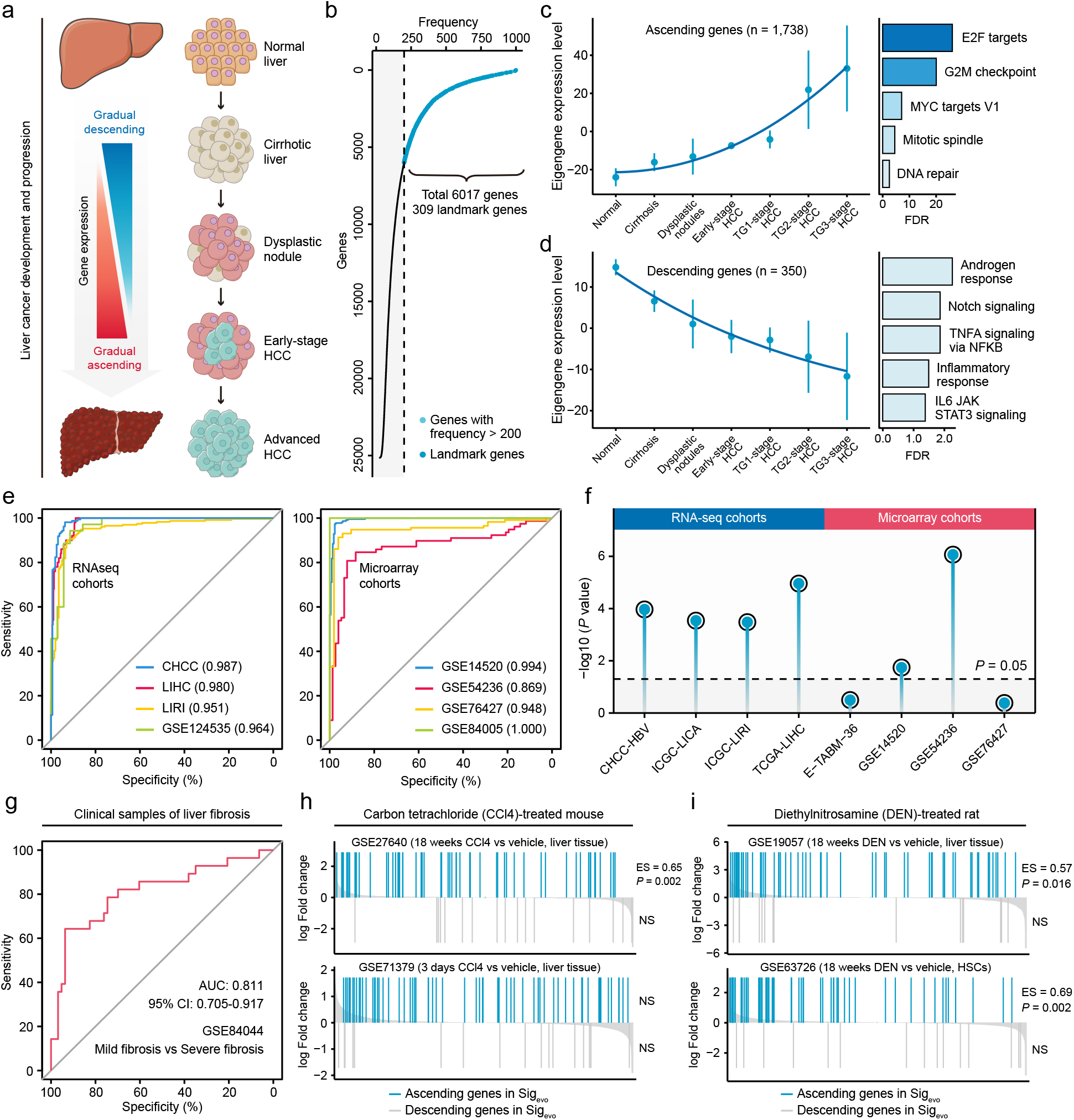
Developing a novel signature, Sig_evo_, for drug repositioning. **a** Schematic of the stepwise process of liver cancer initiation and progression. **b** Preliminary screening of developmental stage-associated genes by Random Forests algorithm. **c** The expression pattern of the ‘ascending’ module discerned by WGCNA analysis (left) and the enriched biological processes determined by hypergeometric test (right). **d** The expression pattern of the ‘descending’ module (left) and the enriched biological processes (right). **e** The performance evaluation of the Sig_evo_ for discerning the difference between tumor and normal tissues based on RNA sequencing cohorts (left) and microarray cohorts (right). **f** The association between the Sig_evo_ and the clinical phenotype of prognosis. Color toward gray indicates no statistical significance. **g** The association between the Sig_evo_ and fibrosis-related phenotype suggested by ROC curve. **h** The association between Sig_evo_ and CCl4-induced expression changes in liver tissues of mice. The enrichment scores and statistical significance were determined by gene set enrichment analysis. **i** The association between Sig_evo_ and DEN-induced expression changes in liver tissues of rats.

As previously discussed, a ‘good’ query signature should reflect the clinical features of corresponding disease comprehensively. To evaluate whether Sig_evo_ was suitable for querying chemopreventive and therapeutic agents for liver cancer, we systematically surveyed the association between this signature and the clinical phenotypes of precancerous/cancerous liver lesions using clinical and experimental data from both human and animal datasets. First, based on clinical cohorts of HCC, we demonstrated that Sig_evo_ had a remarkable capability for distinguishing between tumors from non-tumors, with a median AUC of 0.972 in all eight cohorts (Fig. 4e). Besides, this signature also held great prognostic power in HCC, as suggested by the results of Cox analyses (Fig. 4f). Next, in view of the crucial role of fibrosis in driving hepatocarcinogenesis, further investigation was performed to validate its relevance to fibrosis-related phenotype. The result suggested that Sig_evo_ could also effectively differentiate between mild (S0/S1) and severe (S3/S4) fibrosis, thus demonstrating its relationship with fibrosis progression (Fig. 4g). Additionally, we collected four experimental datasets, including two carbon tetrachloride (CCl4)-treated mouse datasets and two diethylnitrosamine (DEN)-treated rat datasets, to assess the enrichment extent of Sig_evo_ in mouse and rat fibrosis models. It could be observed that ascending genes in Sig_evo_ were significantly enriched in both CCl4-treated (GSE27640) and DEN-treated (GSE19057) liver tissues (Fig. 4 h and i). However, descending genes did not exhibit any significant enrichment pattern in all included datasets, possibly due to the limited gene number (Fig. 4 h and i). Notably, the expression profiles in GSE63726 were derived from non-parenchymal cell fractions which had abundant hepatic stellate cells (HSCs), and thus the significant enrichment could provide the evidence that this signature might reflect the molecular feature of HSC activation (Fig. 4i). In summary, the Sig_evo_ fully complied with the criteria of ‘good’ query signature and was thus employed for querying LINCS.

Subsequently, using the optimal method (XSum) and a compromising parameter (topN = 200), we matched Sig_evo_ with HepG2-derived compound signatures in LINCS and obtained the similarity scores of each compound (lower scores implied higher reversal potency and greater potential for application). After excluding preclinical agents or agents withdrawn from the market according to the drug annotation achieved from the Drug Repurposing Hub, 793 agents remained, which were then considered as repositioning candidates (Additional file 3: Table S11) [24]. Interestingly, some agents which were previously proved to have chemopreventive effects, including erlotinib [25], caffeine [26], and fasudil [27], dominated relatively high rankings on the list (Fig. 5a). Besides, anti-HCC agents were also found to be enriched significantly in compounds with reversal potency (Fig. 5b). These findings converged together to support the reliability of the prediction results.

**Fig. 5.**
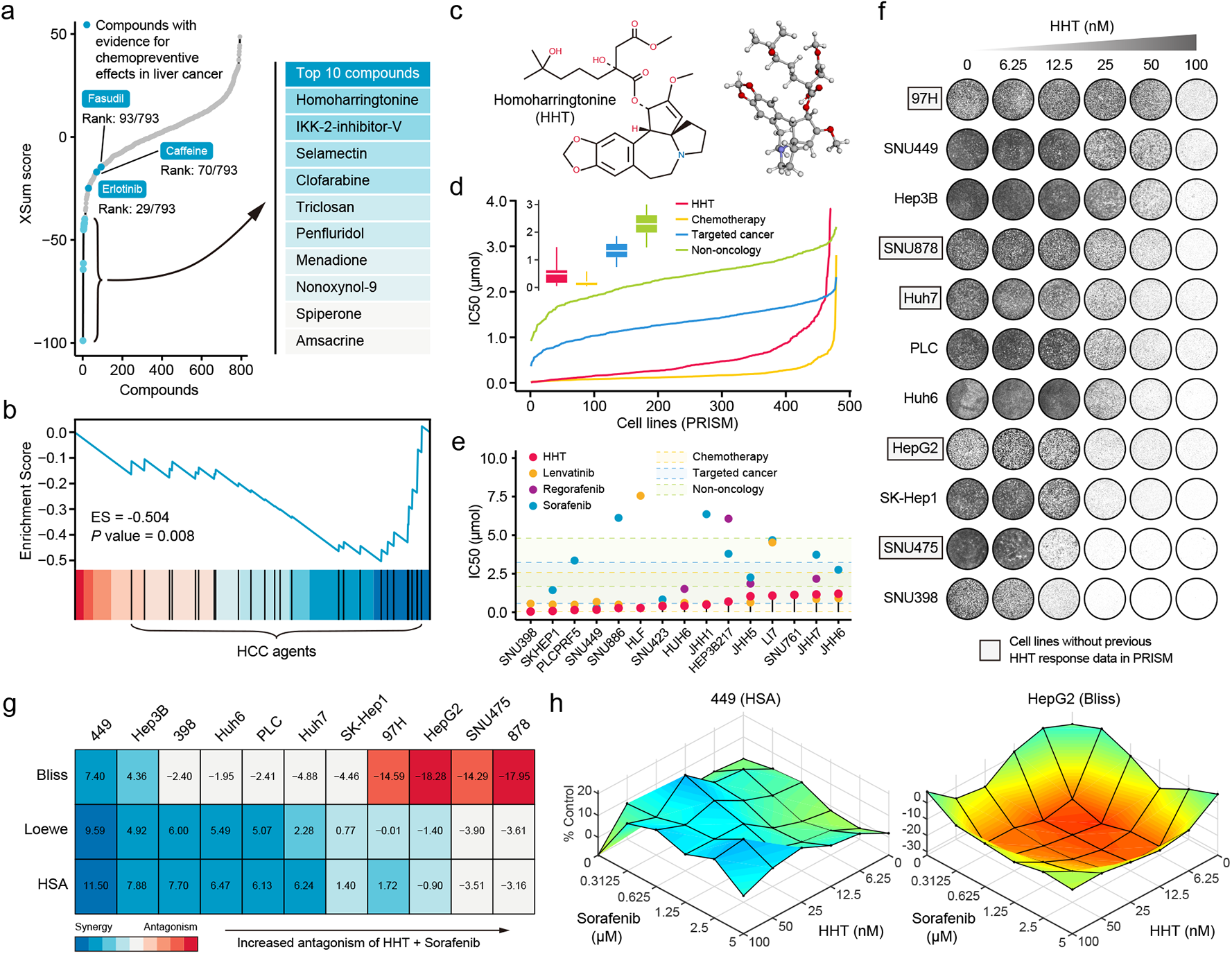
Identifying HHT as repositioning candidate with *in vitro* tumor killing activity. **a** Results of best practice approach-based computational drug repositioning using Sig_evo_ as query signature. Top ranked 10 compounds with highest reversal potency were illustrated in the right panel. **b** Enrichment of HCC agents in compounds with reversal potency (XSum score < 0). The statistical significance was determined based on the null distribution formed by 10,000 permutations. **c** 2D (left) and 3D (right) chemical structure of homoharringtonine (HHT). Data was derived from DrugBank. **d** Comparison of distribution of compound activity between HHT and three different drug categories, including chemotherapy (n = 45 compounds), targeted cancer agents (n = 419 compounds), and non-oncology (n = 362 compounds). The IC_50_ value of each drug category in each cell line (n = 482) was determined through calculating the median IC_50_ value across all the compounds in this category. **e** The drug sensitivity data of HHT (achieved from PRISM dataset) across different HCC cell lines. The drug sensitivities of two HCC agents in the first-line (sorafenib and lenvatinib) and one HCC agent in the second-line (regorafenib) were also illustrated for comparison. Areas with different colors denote the interquartile range of median IC_50_ values of compounds within different drug categories. **f** Long-term cell proliferation assay for testing the anti-tumor activity of HHT across 11 liver cancer cell lines. Of these, five cell lines have not been profiled by PRISM for the sensitivity to HHT. **g** Therapeutic potency of the co-administration of HHT and sorafenib across 11 cell lines. **h** Two representative surface plots showing the overall synergistic (left) and antagonistic (right) effects of the combination treatments.

### 2.8. Characterization of homoharringtonine as anti-liver cancer agent

According to the computational results, homoharringtonine (HHT) (Fig. 5c), a protein synthesis inhibitor targeting RPL3, had the highest reversal potency among 793 repositioning candidates [28]. To ensure the reliability of L1000 platform-derived HHT signature, we treated HepG2 cells with the same condition used in LINCS and utilizing RNA sequencing to measure HHT-induced expression changes. Comparison between L1000-based and RNA sequencing-based HHT signatures was made and a good agreement between them could be observed, suggesting that our prediction using L1000-based HHT signature was comparable and reliable (Additional file 2: Figure S11). Further validation was then conducted to investigate the potency of HHT against liver cancer.

As the drug target of HHT, RPL3 were first characterized and delineated for its clinical and biological implications. The comparison of mRNA expression between normal and tumor tissues suggested that RPL3 exhibited a higher expression in tumor tissues (Additional file 2: Figure S12a). The increase of protein expression of RPL3 could also be observed in tumor tissues, as shown by immunohistochemical images from the Human Protein Atlas (Additional file 2: Figure S12b) [29]. Higher expression of RPL3 also indicated a worse prognosis (Additional file 2: Figure S12c). In addition to the clinical implications, RPL3 also had essential biological implications in liver cancer. Leveraging the clustered regularly interspaced short palindromic repeats (CRISPR)-based screening data from Project Achilles, we found that RPL3 was essential for maintaining the survival and growth of all liver cancer cell lines (Additional file 2: Fig. S12 d and e) [30]. Above results demonstrated the rationality of RPL3 inhibition for treating liver cancer.

Subsequently, we analyzed the drug response data from PRISM which measured the HHT activities in a large panel of cell lines [31]. It could be observed that, compared with molecular-targeted agents and non-oncology agents, HHT had a lower distribution of IC_50_ values across 482 PRISM cell lines (Fig. 5d). Of note, in 15 liver cancer cell lines, HHT has exhibit a powerful anti-cancer capacity with a median IC50 value of 0.408 μM, which was numerically lower than three FDA approved HCC agents (lenvatinib: 0.617 μM; regorafenib: 2.009 μM; sorafenib: 3.348 μM; (Fig. 5e). The anti-liver cancer activity of HHT was also corroborated by the long-term cell proliferation assay (Fig. 5f) and IncuCyte cell proliferation assay (Additional file 2: Figure S13). Given that the co-administration of HHT with other approved agents was more likely to have clinical utility, we also interrogated whether HHT had the ability to augment the tumor-killing effect of sorafenib. Three different statistical models were adopted for synergy estimation. The results suggested that HHT could indeed synergize with sorafenib in many conditions, albeit not very remarkable in general (Fig. 5g-h and Additional file 2: Figure S14). Collectively, our results demonstrated the therapeutic potential of HHT, either administrated as monotherapy or as combination with sorafenib.

### 2.9. Investigation of the anti-fibrotic effects of homoharringtonine

Liver fibrosis occurs when the liver tissue is repeatedly and continuously injured, which is a crucial risk factor for hepatocarcinogenesis [32]. The activation of hepatic stellate cells (HSCs) is one of the key steps in fibrosis development [33]. Since we have proved that Sig_evo_ was associated with liver fibrosis using clinical and animal-derived data, it could be postulated that HHT, a compound with the ability to reverse Sig_evo_, might also be able to inhibit the activation of HSCs and thereby alleviate liver fibrosis. To test this hypothesis, the anti-fibrotic effects of homoharringtonine on TGF-β1-activated human HSC line LX-2 were investigated. We first treated LX-2 cells with vehicle or HHT (1 μM and 5 μM) for 6 hours. Then, RNA sequencing was conducted to quantify the mRNA expression level of vehicle-treated or HHT-treated LX-2 cells (Additional file 2: Table S12). HHT-induced expression changes were obtained based on the resultant data. Nine well-acknowledged fibrotic genes from previous publications were collected for subsequent analysis, the high expression level of which represented the activation status of HSCs. Apparently, the mRNA expression of almost all the fibrosis-associated genes were downregulated after HHT treatments (Fig. 6a and Additional file 2: Figure S15). The downregulated tendency of two most critical genes which encoded collagen I and α-SMA was further corroborated by the quantitative real-time PCR (Fig. 6b). Additionally, the protein-level expression of collagen I and α-SMA was also detected using western blot and immunofluorescence. The results showed that HHT could inhibit the protein expression of collagen I and α-SMA as well (Fig. 6 c and d). Overall, the ability of suppressing HSC activation markers of HHT indicated it had potential anti-fibrotic activity and thus may hold promise for preventing hepatocarcinogenesis.

**Fig. 6.**
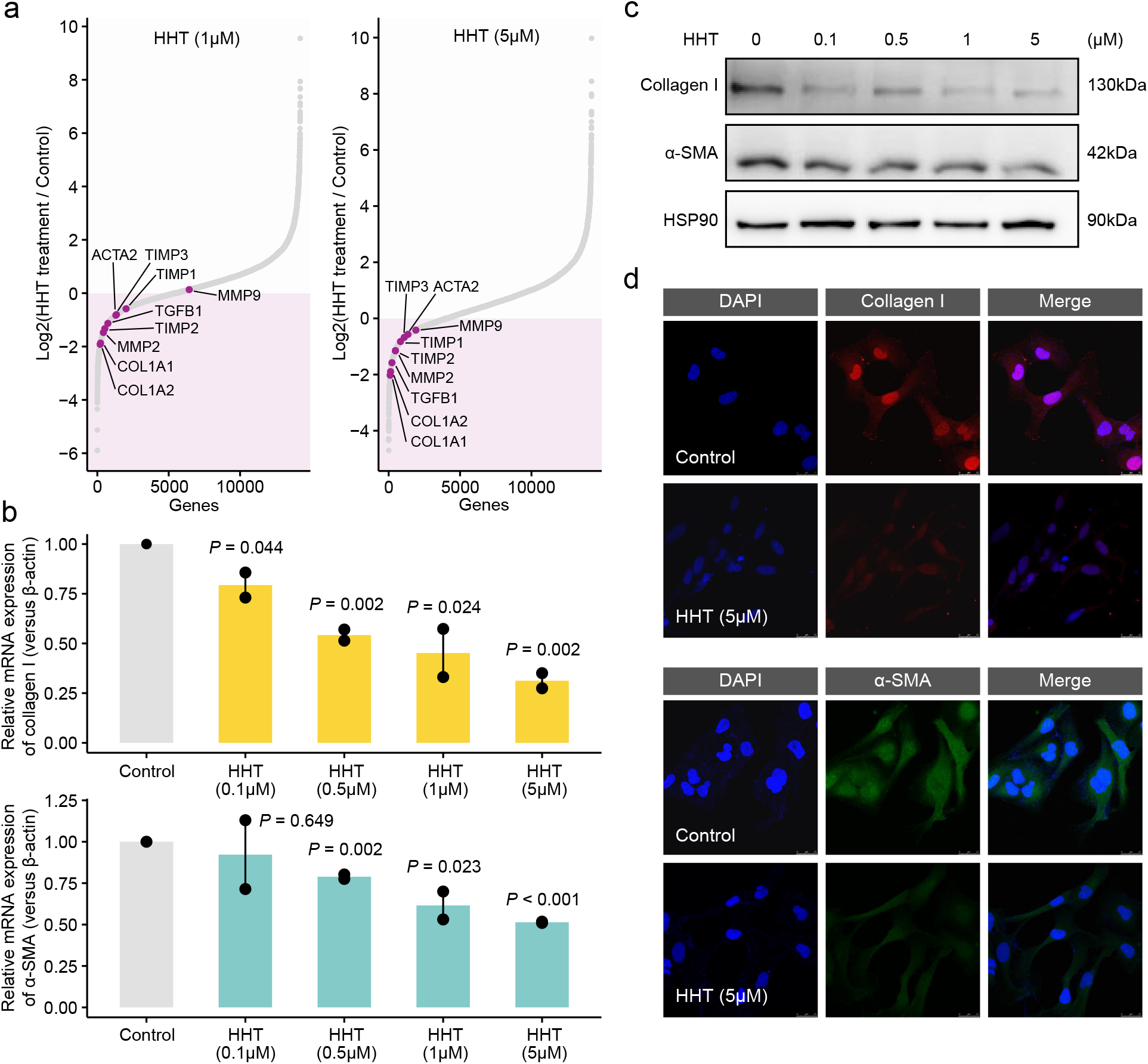
Assessing the anti-fibrotic effects of HHT. **a** Differential expression of nine fibrosis-associated genes between HHT-treated and HHT-untreated LX-2 cells. **b** Quantitative real-time PCR-based mRNA expression level of collagen I (upper) and α-SMA (lower) of LX-2 cells treated with gradient concentrations of HHT for 6 hours. **c** Western blot-based protein expression level of collagen I and α-SMA of LX-2 cells treated with gradient concentrations of HHT for 24 hours. **d** Representative images of immunofluorescence stainings of LX-2 cells with antibody against collagen I (upper) and α-SMA (lower).

## 3. Discussion

In recent years, the explosive growth of pharmacogenomic data enables the development of the new frontier of research named computational drug discovery and repositioning, leading to many remarkable findings of novel therapeutics [34]. Owing to the success of CMap and LINCS projects which profiled numerous compound-induced expression data [4–5], ‘signature reversion’-based computational drug discovery approach has been extensively adopted [7–10]. However, lack of suitable benchmarking standards for evaluating drug repositioning performance limits further improvement of this approach. Although some studies proposed that the benchmarks assessing drug-drug similarity, such as the Anatomical Therapeutic Chemical system (ATC fourth level codes), could be taken as alternative standards to indirectly determine the optimal methodologies and parameters of computational repositioning where gold standards for benchmarking were more complicated [22–23], considering the great difference between these two situations, developing specifically tailored benchmarking standards for assessing disease-drug similarity would be more desirable [21b]. In this study, we proposed two novel benchmarking standards, AUC-based standard and KS statistic-based standard. According to the aforementioned observations, despite being mutually independent, there still existed good agreement of the evaluation results between these two standards, demonstrating the rationality and robustness of the novel standards.

These two standards enable the establishment of a standardized procedure for performing more effective signature-based computational repositioning. We first suggested that using reference signatures from one of the most relevant cell lines with the disease of interest instead of from a non-touchstone cell line or aggregation-based consensus results was a preferable option to exploit LINCS data, in view of a high degree of cell-type specific effects of perturbations. Next, XSum was identified as an optimal method for matching compound and disease signatures. Notably, a prior study which made a comparison of drug retrieval performance between XSum, XCos, and KS methods using a totally different benchmarking standard from ours also come to the same conclusion [21b]. This consistency not only further demonstrated the reliability of the two newly developed benchmarking standards, but also could support our identification of using XSum as the optimal method. Furthermore, we also uncovered the parameter choice (topN = 200) of XSum, which lacked guidance previously.

Within the computational repositioning framework, most of current investigations and methodological developments were focused on reference signatures and signature matching methods. Nevertheless, relatively limited efforts have been made to standardize the procedures for generating quality query signatures [35]. In this study, two potential factors, signature phenotypes and the signature size that might be associated with the retrieval performance of query signatures, were systematically analyzed. First, we intended to figure out how many genes in the query signatures could best characterize a disease state. Through adopting two independent approaches, an appropriate query signature size of 100 was determined. However, some prior studies considered a reduced number of 50 as the optimal length of query signatures [10b, 36]. It is reasonable to speculate that the utility of different signature matching methods (XSum in this study and KS-based methods in other studies) and also the different benchmarking standards may be responsible for the discrepancy. Next, we sought to determine the influences brought by the clinical phenotypes of query signatures, so as to identify the necessary properties a ‘good’ signature needed to possess. Interestingly, the results suggested that a quality query signature should hold the ability to draw a comprehensive picture of clinical features of corresponding disease. This finding seemed to be reasonable since disease was highly likely to be under-represented when creating molecular signature from only one perspective of this disease.

Based on these findings, we summarized the best practice approach for LINCS-based computational drug repositioning. The application of this approach to liver cancer was then carried out. An evolution-associated query signature related to the development and progression of liver cancer was constructed for drug retrieval. Following the best practice approach, HHT was identified as the candidate agent for its highest reversal potency. Since the query signature (Sig_evo_) possessed the liver cancer initiation and development-associated properties simultaneously, we considered that HHT might have both chemotherapeutic and chemopreventive effects for liver cancer accordingly. For validating the chemotherapeutic effect of HHT, drug sensitivity data from PRISM was analyzed and cell proliferation assays were conducted. The remarkable tumor-killing activity of HHT suggested it could be novel therapeutics for liver cancer. For validating the chemopreventive effect of HHT, an indirect approach focused on investigating the anti-fibrotic effect of HHT was adopted. Our results showed that the expression level of fibrosis-associated genes was dramatically decreased when challenging HSCs with HHT treatments. Inhibition of liver fibrogenesis might prevent the progression of cirrhosis and thereby suppress HCC tumorigenesis [25]. Therefore, we supposed that HHT had the potential to be taken as chemopreventive agents for liver cancer as well. Notably, in view of the grim prognosis and imperfect treatment modalities of liver cancer, prevention of HCC development in patients at high risk of primary malignancy or following curative treatment rather than treating patients at advanced stages with limited health benefit is theoretically the most desirable approach to improve patient prognosis [27, 37]. As HHT has been approved by FDA for the treatment of chronic myelogenous leukemia (CML) and widely used in clinical practice without serious adverse effects, it may provide a new chemopreventive strategy that merits further validation in clinical trials [38].

In this study, we have performed the most comprehensive surveys so far about the influencing factors involved in ‘signature reversion’-based drug repositioning. Two novel benchmarking standards are proposed, which provide insight into the evaluation of methodologies developed for signature reversion purpose. Moreover, all the findings in this study are verified independently by at least two different approaches, which can eliminate potential bias and guarantee the reliability of the results. Nevertheless, we also recognize several important limitations. First, with our design, our conclusions are conditional and hold only under the conditions of using compound profiles of HepG2 in LINCS as reference signatures. Further investigations using other LINCS data are required to extend current conclusions to other conditions. Second, the parameters recommended by us, including topN of 200 and query signature size of 100, are more or less based on our subjective judgments and should be taken as a rough guide. Although there are sufficient non-quantitative estimates supporting the use of these two parameters, more efforts are still needed to quantitatively and accurately determine the optimal parameters for this computational approach. Third, in this study we only focused on analyzing the data from the project that utilized transcriptomic platforms to measure cell responses during perturbation experiments, and other omics data which are actively being generated by different LINCS centers might also be a good choice for computational drug discovery and repositioning [39]. Recently, large-scale resources (CPPA) of perturbed protein responses have been generated [40]. Considering that proteins are the components of the basic functional units in biological pathways, investigating the optimal repositioning strategy based on proteomic resources should also have important implication.

## 4. Conclusions

In summary, our findings fill a knowledge gap in the area of LINCS-based computational repositioning. Through exploiting these findings, we also determined a promising anti-liver cancer agent HHT, of which the chemotherapeutic and chemopreventive effects have been validated experimentally.

## 5. Experimental Section

### 5.1. Data collection

LINCS data were downloaded from Gene Expression Omnibus (GEO) database (Phase I: GSE92742, Phase II: GSE70138). Molecular data of clinical and animal datasets were obtained from The Cancer Genome Atlas (TCGA), the Genotype-Tissue Expression (GTEx), GEO and the ArrayExpress databases. Dependency and pharmacogenomic data of cell lines were achieved from the Dependency Map (DepMap) portal. For detailed description, please see the Additional file 1: Supplementary Methods.

### 5.2. AUC-based benchmarking standard

Since establishing benchmarking standards and datasets across all LINCS cell lines seems to be an enormous undertaking, we mainly focused exclusively on one cell line (HepG2) and one disease (liver cancer) in this study. Two benchmarking standards, namely area under curve (AUC)-based standard and Kolmogorov-Smirnov (KS) statistic-based standard, were generated for evaluating the retrieval performance of disease-compound similarity metrics across different conditions. For establishing AUC-based standard, we collected the drug response data from multiple data sources. Compound IC_50_s tested in HepG2 cell line was achieved from ChEMBL (version 27) [41] and Liver Cancer Model Repository (LIMORE) [42] datasets. Compounds among LINCS, ChEMBL, and LIMORE were mapped using compound name followed by manual inspection. Each experiment provided in ChEMBL was also manually checked to ensure the compliance with our requirement. Due to the redundancy of IC_50_s, the median IC_50_s of certain compound among duplicates was used for representing the activity of this compound. We categorized the compounds into effective (IC_50_ < 10μM) and ineffective group (IC50 ≥ 10μM) according to previous study [10b]. The ability to distinguish between effective and ineffective compounds was taken as a measurement of retrieval performance of different similarity metrics (namely AUC value) [10b]. Notably, some have argued that partial AUC (limiting false positive rate at 0.1/0.01) might be a more relevant statistic for actual application of drug repositioning [21b, 22]. However, due to the limited size of our benchmarking dataset, adopting partial AUC with even less information could result in loss of statistical precision, and thus the use of the full AUC is preferred in this situation. The statistical significance of AUCs were calculated through performing permutation test. Briefly, we randomly permuted the class labels of the feature vectors and created 10,000 permutations to form a distribution of ‘random’ AUCs. Then, the *P* value was determined according to the fraction of ‘random’ AUCs greater than or equal to the observed AUC [21b]. For distinguishing, AUC used for evaluating drug retrieval performance was renamed as drug retrieval-associated AUC (DR-AUC), the higher values of which indicate better performance.

### 5.3. KS statistic-based benchmarking standard

To avoid confusion, it should be noted that KS-based method was also used for calculating disease-compound similarity scores, and the specific details were described below. For generating the benchmarking dataset required for the KS statistic-based standard, we systematically surveyed clinical trials involved in HCC treatment and compiled a set of potential HCC agents (clinicaltrials.gov). Preliminary retrieval yielded 1,999 results, and after removing trials failing to fulfil our requirements, we obtained 254 potential therapeutic agents for HCC. The detailed retrieval process was presented in Figure 3A. To minimize potential selection bias, this process was performed independently by two investigators (C.Y. and X.H.). Perturbagen-induced expression profiles of 27 agents among these tested in HepG2 cell line are available in LINCS dataset. Based on the assumption that HCC-associated agents are more likely to reverse HCC signature than random agent combinations, the enrichment capabilities of different similarity metrics could be used to assess their repositioning performance. The calculation of enrichment score (ES) of HCC agents was generally identical with that in gene set enrichment analysis [43]. Considering that there was no need accounting for the size of the agents set, we did not calculate the normalized enrichment score (NES) that might introduce additional randomization. To obtain a nominal *P* value, we created 10,000 permutations and recomputed the ES for each permutation to form a null distribution. The significance *P* of the observed ES was then determined relative to the null distribution. Notably, ES used in retrieval performance evaluation was renamed as drug retrieval-associated ES (DR-ES), the lower values of which represent better performance.

### 5.4. Signature matching methods

The retrieval performance of six different matching methods, including eXtreme Sum (XSum) [21b], eXtreme Cosine (XCos) [21b, 22], eXtreme Pearson (XCor) [23], eXtreme Spearman (XSpe) [23], Kolmogorov-Smirnov (KS) test [4], and the Reverse Gene Expression Score (RGES) [10b], was systematically benchmarked. Based on the consideration that small variations in expression changes may be noise without biological significance, the eXtreme methods only utilized top up- and down-regulated genes in compound signatures for similarity score calculation (all remaining genes were assigned the values of zero). By comparison, KS and RGES methods use complete compound profiles as reference signatures. The detailed features and scoring schemes of these methods are described as follows.

The XSum method handles the up- and down-regulated genes separately. In brief, the sums of the change values in reference/compound signatures relative to up-regulated query/disease genes (sum_up_) and down-regulated query/disease genes (sum_down_) are first calculated. Then, XSum is defined as following: XSum = sum_up_ - sum_down_. Other three eXtreme methods, including XCos, XCor, and XSpe, take disease signatures as a whole to query compound signatures, and they calculated the correlation between the numeric vectors of disease and compound signatures using cosine similarity, Pearson correlation, and spearman correlation, respectively. Notably, cosine similarity is nearly identical with Pearson correlation except without centering vectors around the mean values. The KS method was adopted by the first CMap study and has been the most widely used method for connecting disease signatures to compound signatures [4]. Similar to XSum, KS method also needs to sperate disease signatures into two gene sets, up-regulated gene set and down-regulated gene set, and ignores the magnitude of differential expression. Briefly, using complete compound profiles as reference, maximum deviation (MD)-based enrichment scores of up-regulated gene set (es_up_) and down-regulated gene set (es_down_) are first computed. If es_up_ and es_down_ have the same algebraic sign then KSscore = 0, otherwise, KSscore = es_up_ - es_down_. The RGES method is a recently proposed modification of the original KS method, which was demonstrated to perform better in drug response prediction than KS method [10b]. In contrast to original KS method, RGES focuses on the reversal relation between the disease and agents, and RGES is defined as es_up_ - es_down_ regardless of the sign of es_up_ and es_down_. In addition to the above six methods, there also exist many other methods, such as WSS/sscMap [44], TES [45], ProbCmap [36], NFFinder [46], EMUDRA [23], and so on, for calculating the similarity between disease and compound signatures. However, some of them are not accessible currently and some are developed based on the data of initial CMap dataset (1,309 compounds), and accordingly, we did not include these methods into our analyses.

### 5.5. Generation of query signatures for performance evaluation

For evaluating retrieval performance of similarity metrics at different conditions, we prepared four HCC-associated gene signatures to query LINCS. Two of them, Sig_gastro_ and Sig_NC_, are achieved from previously published studies [10]. Given that the development of these two signatures were mainly based on LIHC cohort, as a complement, the other two were generated from another RNAseq cohort (LIRI) and a microarray cohort (GSE54236), respectively. The differentially expressed genes in LIRI cohort were computed using *edgeR* package (version 3.26.5) on raw count data [47]. For microarray data, we used *limma* package (version 3.40.2) to conduct differential expression analysis on normalized data [48]. The statistically significant differential genes in above analyses were defined as adjusted *P* < 0.01 and absolute log_2_ fold change (FC) > 1. As a result, we obtained a 70-gene signature (Sig_LIRI_) with 48 up- and 22 down-regulated genes from LIRI and a 28-gene signature (Sig_GSE54236_) with 22 up- and 6 down-regulated genes from GSE54236 respectively, which could represent discordant expression pattern of HCC. Considering that the gene numbers in signatures created through differential expression analysis were much less than that in prognostic signatures (see section below), to make comparison of retrieval performance between these two signature phenotypes without being subject to the signature size, we relaxed the significance threshold of differential genes to *P* < 0.01 and log_2_FC > 0.5, and built two increased signatures which included 125 genes (LIRI) and 116 genes (GSE54236) respectively. These two increased signatures were also used to explore the potential influences of signature size.

### 5.6. Construction of size-diversified query signatures

We adopted two independent approaches to explore whether the differences of query/disease signature size could affect subsequent drug retrieval. The first approach was based on iterating the threshold of fold change values, ranging from 0.1/-0.1 to the maximum/minimum with a increment/decrement of 0.05, which could obtain a number of query signatures with a diversity of signature size (duplicates were removed). As for the second approach, above two increased signatures, 125-gene signature from LIRI and 116-gene signature from GSE54236, were taken as the basis for generating smaller-size testing signatures. Briefly, we randomly extracted testing signatures from complete signatures, with the size ranging from the minimum of five to the maximum of 124 or 115. To avoid bias, this process was repeated 1,000 times to generate 1,000 testing signatures for each signature size.

### 5.7. Construction of query signatures representing different clinical phenotypes

To investigate whether the clinical phenotype of signature was potential factor affecting the retrieval performance, we developed two strategies, a forward strategy starting from generation of signatures with distinguishing clinical phenotypes to the evaluation of retrieval performance and a backward strategy starting from obtaining signature with the best performance to the comprehensive investigations of its clinical implication. For the first strategy, to compare with general HCC signature representing discordant expression pattern, two prognostic signatures based on LIRI and GSE54236 cohorts were constructed. We integrated survival data with expression data and performed Cox proportional hazards regression to assess association between overall survival and gene expression. The statistically significant prognostic genes were defined as *P* < 0.005. A 133-gene prognostic signature with 117 poor- and 16 good-outcome genes was generated based on LIRI, while analysis on GSE54236 resulted in a 107-gene prognostic signature with 79 poor- and 28 good-outcome genes. Comparisons of drug retrieval performance between these two types of signatures were carried out subsequently. For the second strategy, taking 978 landmark genes as a basis, simple random sampling without replacement (SRSWOR) was performed to extract genes from landmarks for forming candidate signatures. The size of randomized signatures was set at 100 and the process of random sampling was repeated 10,000 times to obtain a collection of 10,000 randomized signatures. The DR-AUC and DR-ES values were then calculated for each signature, and the optimal one was defined as the signature with the minimum of DR-AUC multiplying DR-ES.

### 5.8. Generation of evolution-associated query signature

To find compounds with potential in the prevention and treatment of liver cancer, we developed a hepatocarcinogenesis and progression-associated signature. GSE89377 cohort was utilized to build this signature while GSE6764 cohort was taken for external validation. To identify genes associated with developmental stages, we constructed Random Forests (RF) model taking stages as dependent variable. Variable importance was assessed with the mean decrease accuracy (MDA) measures for individual factors in RF model. Variables with positive MDA values are of high importance in predicting stages. In other words, these variables are more likely to be related with liver cancer development and progression (negative MDA values can be regarded as equivalent to zero importance with no predictive power). The RF analysis was independently repeated 1,000 times with 1,000 trees growing each time, and genes with positive MDA values incorporated in more than 200 iterations were kept for subsequent analyses.

We next performed weighted gene co-expression network analysis (WGCNA) to assign resultant genes into modules according to expression similarity, and simultaneously, to recognize the trajectories of gene expression during liver cancer development [49]. First, an appropriate soft threshold was estimated by using the *pickSoftThreshold* function in *WGCNA* package. Then, we constructed WGCNA network and detected gene expression modules using *blockwiseModules* function with a minimum module gene number of 50, soft thresholded power of 12, and a dendrogram cut height of 0.3. Genes without assignment to specific modules were assigned the color of grey. Module eigengenes (MEs) representing the first principal components (PC1) of each module were returned, and the module-trait relationship (MTR) analysis was conducted by calculating the correlation between MEs and developmental stages. The expression trend of each module across seven stages of HCC development was visualized through using mean PC1 values of samples in each stage to generate trend curves. According to the correlation coefficient of MTR analysis and the visualized expression trend of each module, two modules exhibiting the highest positive/negative correlation with developmental stages as well as showing gradually increasing or decreasing expression trends were selected for their potential functional roles during HCC initiation and development. Subsequently, to explore the biological processes associated with genes in these two modules, we conducted hypergeometric test based on the hallmark gene sets (h.all.v7.0.symbols) downloaded from the Molecular Signatures Database (MSigDB) using *enricher* function in *clusterProfiler* package [50]. The *P* values from the hypergeometric tests were adjusted for multiple comparison testing and an adjusted *P* value less than 0.05 was considered significantly enriched.

Genes in these two modules were mapped to the 978 landmark genes, resulting in a 159-gene panel (134 genes in ascending module and 25 genes in descending module). According to the findings described in *Results* section, we decided to further narrow down this panel to create a query signature with the size of 100 genes. The molecular and clinical data in GSE15654 were utilized to determine the association between the expression patterns of signatures and the occurrence of HCC. Briefly, we first performed SRSWOR to extract a subset of 100 genes from the 159-gene panel, repeated 10,000 times. The processes of random sampling were conducted separately for ascending and descending genes, to ensure a minimum descending gene number of two in the randomized signatures. As a result, 10,000 randomized signatures with 100 genes per signature were generated. Next, PC1 values of all randomized signatures were extracted based on expression data from GSE15654 to represent the overall expression patterns of these signatures, and the follow-up data using HCC occurrence as endpoint was then integrated with above expression pattern data for subsequent Cox proportional hazards regression (COXPH). The signature which had the minimum *P* value across the 10,000 COXPH analyses was considered as the optimal signature and retained for querying LINCS. The expression trend of this signature was further validated GSE6764 cohort.

### 5.9. Experimental methods

All experimental methods, including cell proliferation assays, quantitative real-time PCR, western blotting analysis, immunofluorescence, RNA sequencing, etc., were described in the Additional file 1: Supplementary Methods.

### 5.10. Statistical analysis

All the computational analyses and graphical visualization were performed in R software, version 3.6.0 (https://cran.r-project.org/). Unless stated otherwise, correlation between two continuous variables was measured by Spearman’s rank-order correlation, and pairwise comparisons were conducted using Kruskal-Wallis and Wilcoxon sum rank tests. ROC curves and AUC values were visualized and calculated using the pROC package[51]. The hazard ratio (HR) was estimated using Cox regression model in *survival* R package. Cumulative hazard curve was carried out using *jskm* package and the log-rank test was used to determine the statistical significance of differences. A two-tailed *P* < 0.05 was considered significant unless indicated otherwise.

## Additional files

Additional file 1: Supplementary methods and Supplementary discussion. (DOCX 113 kb)

Additional file 2: Supplementary figures 1-15. (PDF 4333 kb)

Additional file 3: Supplementary tables 1-12. (XLS 2279 kb)

## Availability of data and materials

Sequencing data necessary for the analysis are available within Supplementary Data. The rest of the data supporting the findings of this study are available from the corresponding author upon reasonable request. The codes are available at https://github.com/YangJAT/Best_practices_for_LINCS.

## Acknowledgements

This work was financially supported by the National Natural Science Foundation of China (81972208), Shanghai Natural Science Foundation (19ZR1452700).

## Conflict of interest

The authors declare no conflict of interest.

## Authors’ Contributions

C.Y. and M.C. designed the study and performed the computational analyses. S.W. and X.H. and R.Q. performed the *in vitro* validation experiments. C.Y., J.W. and Z.L. wrote the manuscript. Q.W. provided guidance on manuscript preparation. H.W., H.H. and C.W. supervised the study.

## Notes

### Competing Interest Statement

The authors have declared no competing interest.

