## Additional file 1 for "A survey of optimal strategy for signature-based drug repositioning and an application to liver cancer"

### Supplementary Methods

#### LINCS data source and processing

We downloaded the LINCS level 5 data (moderated Z-score) which comprising the differential expression signatures for nearly 20,000 unique compounds as well as meta-information of these signatures from Gene Expression Omnibus (GEO) database (Phase I: GSE92742, Phase II: GSE70138). Because this study only focused on analyzing compound signatures, those signatures induced by other perturbagens including gene knockdown/knockout and gene overexpression were first excluded. The L1000 platform used by LINCS project only measures the expression level of 978 landmark genes, and the expression of remaining genes was based on imputation [1]. This set of landmark genes are widely expressed in various cellular contexts and can well represent the full genome [1]. Accordingly, we chose to use just the landmark genes [2, 3]. To ensure the reliability, only high-quality signatures are designated for following analyses (is_gold=1, see https://clue.io/connectopedia/glossary for detailed information). In addition, L1000 data was further filtered for only 6-hour treatment samples due to the most abundant experiments on HepG2 cell line in this time point. Besides, previous study also showed that gene expression changes obtained at a late time point (such as 24h) might reflect secondary or even tertiary responses, and the mechanistic effects of compounds might not be correctly recorded at late time point [4]. As for the perturbation concentration, we selected expression profiles measured at 10μM considering that this relatively high concentration is often chosen for performing high-throughput small molecular screens, and also, there exist the most abundant experiments at this concentration. The similarity between compound pairs was calculated based on cosine similarity algorithm [5]. For visualizing the LINCS data in 2D space, we measured the cosine distance (1-cosine similarity) between signatures and utilizing cosine distance matrix as input to perform t-distributed stochastic neighbor embedding (t-SNE) analysis [6]. We downloaded the mechanism of action (MOA) and clinical phase information of compounds in LINCS from the Drug Repurposing Hub (https://clue.io/repurposing) [7]. The basal expression data of LINCS cell lines was achieved from the Broad Institute Cancer Cell Line Encyclopedia (CCLE) project (https://portals.broadinstitute.org/ccle/) [8]. The expression-based similarity between different cell lines or between cell lines and clinical samples was determined through using ranked-based Spearman correlation [3].

#### Other pharmacogenomic datasets

In addition to ChEMBL and LIMORE, the drug response data of HepG2 cell line was also obtained from the Cancer Therapeutics Response Portal (CTRP) dataset (CTRPv.2.0, released October 2015) [9]. Considering that $IC_{50}$s were not provided, the available area under the dose-response curve (AUDRC) values were used solely for demonstrating the correlation between reversal potency and drug efficacy in different conditions. The AUC values in CTRP range from 0 to 30, and similar to $IC_{50}$, lower values indicate increased sensitivity to treatment. To investigate the drug sensitivity of repositioning candidate across different HCC cell lines, we achieved the response data from the PRISM Repurposing dataset (19Q4, released December 2019). Although $IC_{50}$s are also provided by PRISM as one of drug response metrics, the drug response data of HepG2 cell line is absent (similar situation also exists in the Genomics of Drug Sensitivity in Cancer dataset). Therefore, drug response data in these datasets were not used for developing AUC-based benchmarking standard. A summary of included pharmacogenomic datasets was presented in Table S1.

**Genetic Dependency Data**

CRISPR dependency data were obtained from the 20Q1 dependency map (DepMap) portal, which contained dependencies estimated for nearly 20,000 protein coding genes and 739 cell lines using the CERES algorithm [10]. CERES score (gene effect) was used to measure the dependency of the gene of interest in cell lines, and a lower CERES score indicates a higher likelihood that the gene is essential in cell growth and survival. Besides, data of dependency probability were also achieved. A probability of dependency of certain gene in certain cell lines greater than 0.5 represents that the gene can be considered essential in this cell line. Essential genes in liver cancer were defined as genes that were essential in all 22 liver cancer cell lines.

**RNA-sequencing datasets**

We collected five RNA-sequencing (RNAseq)-based HCC cohorts, including CHCC-HBV [11], LICA-FR [12], LIRI-JP [13], TCGA-LIHC [14], and GSE124535 [15], representing 947 HCC patients derived from four geographically different origins. Of these, LICA, LIRI, and LIHC cohorts provided raw counts quantifying gene expression, which were transformed into transcripts per kilobase million (TPM) values for subsequent analyses (raw counts were only used for *edgeR*-based differential expression analysis) [16]. CHCC and GSE124535 cohorts provided fragments per kilobase per million reads (FPKM) normalized data, which was also converted to TPM values. All TPM values were log2 transformed. In addition, the batch effects-normalized expression matrices of ~ 10,000 patients across 33 human cancers (TCGA Pan-Cancer) were downloaded from the UCSC Xena browser (http://xena.ucsc.edu/). The RNAseq expression profiles of 29 normal tissues were downloaded from the Genotype-Tissue Expression (GTEx) project (https://gtexportal.org/home/). Ensembl GeneIDs were mapped to HGNC symbols using *biomaRt* package.

**Array datasets**

Five microarray-based clinical cohorts, including E-TABM-36 [17], GSE14520 [18], GSE54236 [19], GSE76427 [20], and GSE84005, were included to construct and validate HCC-associated signatures. Raw microarray data generated from Affymetrix platforms were normalized using robust multi-array average (RMA) method in *Affy* package [21], while Illumina platform-derived raw data were normalized using the robust spline normalization (RSN) method in *lumi* package [22]. In other cases, normalized data was directly downloaded for use. Three liver cancer development-associated cohorts, including GSE89377, GSE6764 [23], and GSE15654 [24], were also included for constructing the new signature, $Sig_{evo}$, which was then applied to query LINCS for finding potential therapeutics of liver cancer. The samples in GSE89377 and GSE6764 covered multiple stages of the development of liver cancer, with the ability to recapitulate the stepwise process of hepatocarcinogenesis and progression. As for GSE15654, this cohort contains the gene expression profiles of samples from 216 patients with early cirrhosis who were prospectively followed for a median of 10 years, which thus can be used to identify the relationship between gene expression and HCC occurrence [24]. In addition, a liver fibrosis-associated clinical cohort (GSE84044) [25] and four experimental datasets, including two carbon tetrachloride (CCl4)-treated mouse datasets (GSE27640 and GSE71379) [26, 27] and two diethylnitrosamine (DEN)-treated rat datasets (GSE19057 and GSE63726) [26, 27], were utilized to further assess the potential implication of $Sig_{evo}$. Mouse and rat genes were mapped to orthologous human genes using *biomaRt* package, and genes without known human homologous relationships were excluded. A summary of detailed information of all RNAseq and array datasets has been presented in Supplementary Table 2.

**Clinical data**

Among the clinical cohorts above, nine cohorts have corresponding follow-up information, including four RNAseq cohorts (CHCC, LICA, LIRI, and LIHC), and five microarray cohorts (E-TABM-36, GSE14520, GSE54236, GSE76427, and GSE15654). For RNAseq cohorts, the survival data of CHCC and LICA cohort was obtained from the supplementary files of reference [11, 12], data of LIRI cohort was achieved from the International Cancer Genome Consortium (ICGC) portal (https://dcc.icgc.org/), and data of LIHC cohort was achieved from TCGA Pan-Cancer Clinical Data Resource (TCGA-CDR). For microarray cohorts, complete clinical data was accessed from either public database (GEO: https://www.ncbi.nlm.nih.gov/gds/; ArrayExpress: https://www.ebi.ac.uk/arrayexpress/) or the original authors. Notably, except GSE15654 which uses the occurrence of HCC as endpoint, other eight cohorts all take survival status as endpoint.

**Human cell lines and compounds**

The liver cancer cell lines, Hep3B, Huh7, PLC/PRF/5, SNU398, Huh6 were provided by Erasmus University (Rotterdam, Netherlands). MHCC97H and SK-Hep1 were provided by the Liver Cancer Institute of Zhongshan Hospital (Shanghai, China). SNU449, SNU878, SNU475 and HepG2 was purchased from the American Type Culture Collection (ATCC). The immortalized human hepatic stellate cell (HSC) line LX2 was also purchased from the ATCC. These cells were maintained in Dulbecco's modified Eagle's medium (DMEM) (Gibco, Carlsbad, CA) supplemented with 10% fetal bovine serum (FBS) (Gibco) and 1% penicillin/streptomycin (BasalMedia), incubated at 37°C in humidified atmosphere with 5% CO2. Mycoplasma contamination was excluded via a PCR-based method. The identities of all the cell lines were confirmed by short tandem repeat (STR) profiling. Human recombinant transforming growth factor β1 (TGF-β1) was purchased from R&D Systems (Minneapolis, MN, USA), which was used to activate LX2 (10 ng/mL TGF-β1 for 24h). HHT treatment was performed by pre-treating for 2 h before TGF-β1 stimuli. Homoharringtonine (S9015) was purchased from Selleck Chemicals and dissolved in dimethyl sulfoxide (DMSO) using a storage concentration of 10mM.

**Cell proliferation assays**

For long-term cell proliferation assay, cells were seeded into six-well plates (2-3×$10^4$ cells per well) and HHT was added after 24 hours. Cells were treated with HHT as indicated for 10 days during which the culture media were replaced every three days. Then, cells were stained with 1% crystal violet for 10 minutes and rinsed with tap water. Pictures were taken using ImageScanner$^{TM}$ III (GE Healthcare) at 300 dpi resolution. For IncuCyte cell proliferation assay, cells were cultured and seeded into 96-well plates at a density of 1000–1500 cells per well, and 24 hours later, HHT was added at indicated concentrations. Cells were imaged every 4 hours in IncuCyte ZOOM system (Essen Bioscience) and phase-contrast images were collected and analyzed to determine the proliferation curves based on cell confluence.

**Quantitative real-time PCR**

We first harvested cells using TRIzol reagent (Invitrogen) based on manufacturer's instruction. Then, cDNA synthesis was carried out using Maxima Universal First Strand cDNA Synthesis Kit (no.K1661, Thermo Scientific). Quantitative reverse transcription PCR (qRT-PCR) assays were conducted using 7500 Fast RealTime PCR System (Applied Biosystems). Relative mRNA levels of genes shown were normalized to the mRNA level of glyceraldehyde-3-phosphate dehydrogenase (GAPDH) (housekeeping gene). The primer sequences for assays using SYBR Green master mix (Roche) are as follows: β-actin Forward, AAATCTGGCACCACACCTTC; β-actin Reverse, GGGGTGTTGAAGGTCTCAAA; Collagen I Forward, TCCTGGTCCTGCTGGCAAAGAA; Collagen I Reverse, CACGCTGTCCAGCAATACCTTGA; α-SMA Forward, GACAATGGCTCTGGGCTCTGTAA; α-SMA Reverse, CTGTGCTTCGTCACCCACGTA.

**Western blotting analysis**

Cells were washed with PBS and lysed on ice with RIPA lysis buffer supplemented with Complete Protease Inhibitor (Roche) and Phosphatase Inhibitor Cocktails II and III (Sigma). Protein concentration was measured using the BCA Protein Assay Kit (Pierce). All lysates were then freshly prepared and processed with Novex NuPAGE Gel Electrophoresis Systems (Thermo Fisher Scientific) followed by western blotting. The antibody against α-smooth muscle actin (α-SMA) (A5228) was obtained from Sigma-aldrich (USA) and the antibody against collagen I (14695-1-AP) was achieved from Proteintech.

**Immunofluorescence**

Cells were cultured on glass cover slips, fixed for 10 minutes with 4% formaldehyde, and permeabilized with 0.5% Triton X-100 for 15 minutes at room temperature. Immunofluorescence analysis was performed using the following antibodies: Anti-Actin, α-Smooth Muscle antibody (10μg/ml), Anti-collagen I (1:1000), Anti-mouse IgG Fab2 Alexa Fluor (R) 488 (1:2000, CST), and Anti-rabbit IgG Fab2 Alexa Fluor (R) 542 (1:2000, CST). Cell nuclei were stained with DAPI (4,6-diamidino-2-phenylindole). After immunostaining, the samples were observed using a LEICA TCS SP5 confocal microscope.

**RNA sequencing**

For RNA sequencing, total RNA was extracted and purified using the Trizol reagent (Invitrogen). The library was prepared using TruSeq RNA sample prep kit according to the manufacturer's protocol (Illumina). Paired-end libraries were sequenced by an Illumina HiSeq X Ten (2 x 150-nucleotide read length), with a sequence coverage of 20 million paired reads. For data analysis, raw sequencing reads were mapped to the human genome (GRCh38) using STAR (version 2.4.2g1). Then gene-level read counts were generated using featureCounts from the subRead package with default settings.

### Supplementary Discussion

We summarized several potential factors that might affected computational drug repositioning. Selecting appropriate compound signature (also termed reference signature) is the first major challenge encountered during the process of matching disease and compound. Four influencing factors associated with compound signatures were considered in this study:

**Raw data processing:** Raw scan data accessed from Luminex FlexMap 3D scanners must be deconvoluted from only 500 Luminex bead colors to obtain the expression level of each gene in landmark panel. The initial LINCS project applied the k-means clustering algorithm for handling this process [1]. Recently, some alternative methods, such as Gaussian mixture model (GMM)-based and Bayesian-based methods, have been proposed for performing peak deconvolution, which might measure the expression changes more accurately, thereby improving the performance of down-stream drug discovery and repositioning [28, 29].

**Source of cell line:** Previous studies have pointed out that a compound might induce inconsistent transcriptional changes across different cell lines, indicating that using signatures generated from unrepresentative cell line or using average/consensus signatures merged from multiple different cell lines could lead to incorrect down-stream conclusions [1, 30, 31]. In the present study, we also systematically surveyed this factor and demonstrated the presence of a high degree of cell-specific perturbed expression responses. According to these findings, researchers should consider the relevance of the cell lines used in LINCS to the diseases on which they are focused, and obtain signatures based on one of the most relevant cell lines for subsequent drug retrieval.

**Perturbation time and dose:** Currently, no consensus regarding the most appropriate perturbation time for connectivity mapping-based drug discovery and repositioning has been achieved. The initial CMap study selected a relatively early time point (6 hours) after compound treatment for recording transcriptomic changes related to direct mechanisms of action, since profiles obtained at a late time were considered to reflect secondary and tertiary responses [32]. Besides, another study also mentioned that longer duration of perturbation (such as 24 hours) might lead to phenotypic changes unrelated to the perturbagens rather than mechanistic changes [4]. However, a previous study made a comparison between 6-hour and 24-hour treatment data, and found that there was no significant difference in the performance of target prediction [33]. As for concentration, considering that the optimal perturbation concentration is not known for many compounds of potential interest, the initial CMap adopted a relatively high concentration of 10μM to perform perturbation experiments [32]. Besides, using this relatively high concentration also ensures that perturbations can reach the minimum criteria causing a measurable transcriptomic changes. To date, due to many limitations, there are still no studies investigating which perturbation time and dose can yield best drug retrieval performance. In this study, all the analyses involved in LINCS were performed using data from experiments based on a fixed perturbation time of 6 hours and concentration of 10μM, and the issues about the optimal time and dose selection remained unaddressed.

As with compound signature, disease signature (also termed query signature) might also affect the process of pattern matching. Three potential factors influencing the generation of disease signatures were discussed and investigated:

**Quality of dataset:** The quality of clinical datasets used to create the disease signatures directly determine the representativeness and reliability of these signatures. Many factors during collecting and analyzing clinical samples can affect the quality of the resultant datasets. For example, sampling bias caused by unskilled sampling operators or highly heterogeneous tumors may lead to the limited representativeness of collected samples, and disease signatures based on these samples are unlikely to recapitulate the actual features of corresponding diseases. In addition, another potential factor that cannot be ignored is the detection limits of expression profiling technologies, and in our experience, the expression measurements of some old platforms are not particularly accurate, which may also affect the reliability of resultant signatures. Generally, if possible, generating disease signatures using datasets previously proved to be of high quality and reliable seems to be a sound approach to mitigate above concerns. In addition, integrating gene signatures from diverse studies with similar settings may also be an alternative method which also has the potential to generate robust and stable signature representation [34].

**Clinical phenotype of the signature:** Through integrating molecular profiles with various clinical data, signatures representing different phenotypes can be achieved, such as differential expression-associated signature (adopted by most of studies), prognosis-associated signature [35], and metastasis-associated signature [36]. Currently, there are limited studies exploring the potential effect of clinical phenotype on drug retrieval. A previous study made a comparison of drug prediction performance between a general breast cancer signature and a validated breast cancer prognostic signature, and found that the querying results based on the prognostic signature were more likely to have clinical significance [35]. However, considering that it was a case study without controlling potential bias caused by dataset and signature size, more efforts are required to achieve a more robust conclusion.

**Number of genes in the signature:** Query signature size is another potential factor affecting the accuracy of disease-compound matching. The choices of the signature length vary drastically from one study to another in real applications, ranging from tens to thousands [37, 38]. Using the full-genome data from the original CMap dataset, top (respectively, bottom) 250 genes were identified as the components of optimal signatures [30]. When using CMap data based on a decreased panel of landmark genes, 50 genes were considered as an optimal signature size [39]. Similarly, exploiting the landmark gene expression data in LINCS, another study pointed out that the size of disease signatures should be more than 50 genes to achieve best performance [2]. Despite these findings, it is worth noting that above studies all utilized KS statistics-based methods for measuring connectivity mapping when investigating this factor, and the optimal signature size for conducting eXtreme-based methods has still been under-explored.

Once both compound signatures and disease signatures are ready, the last and also the most important step is to calculate similarity scores between two signatures, so as to identify potential therapeutic agents for corresponding diseases.

**Disease-compound matching algorithms for drug retrieval:** The initial CMap study utilized KS statistic to connect disease to compound [32]. After that, many alternative similarity methods have been developed. However, due to the lack of gold standards for evaluating disease-compound matching performance, the comparisons of the accuracy of drug prediction between these methods have always been considered as a big challenge. Fortunately, there exist relatively clear standards for assessing drug relationship prediction, and the performance of these methods can thus be determined. Current available similarity algorithms include methods developed from the standard KS statistic, such as TES [30] and RGES [2], methods based on eXtreme theory, such as XSum [35], XCos [5, 35], XCor [40] and XSpe [40], and other types of methods, such as WSS/sscMap [41, 42], ProbCmap [39], NFFinder

[43], and EMUDRA [40].
