## Additional file 2 for "A survey of optimal strategy for signature-based drug repositioning and an application to liver cancer"

### Supplementary Figures

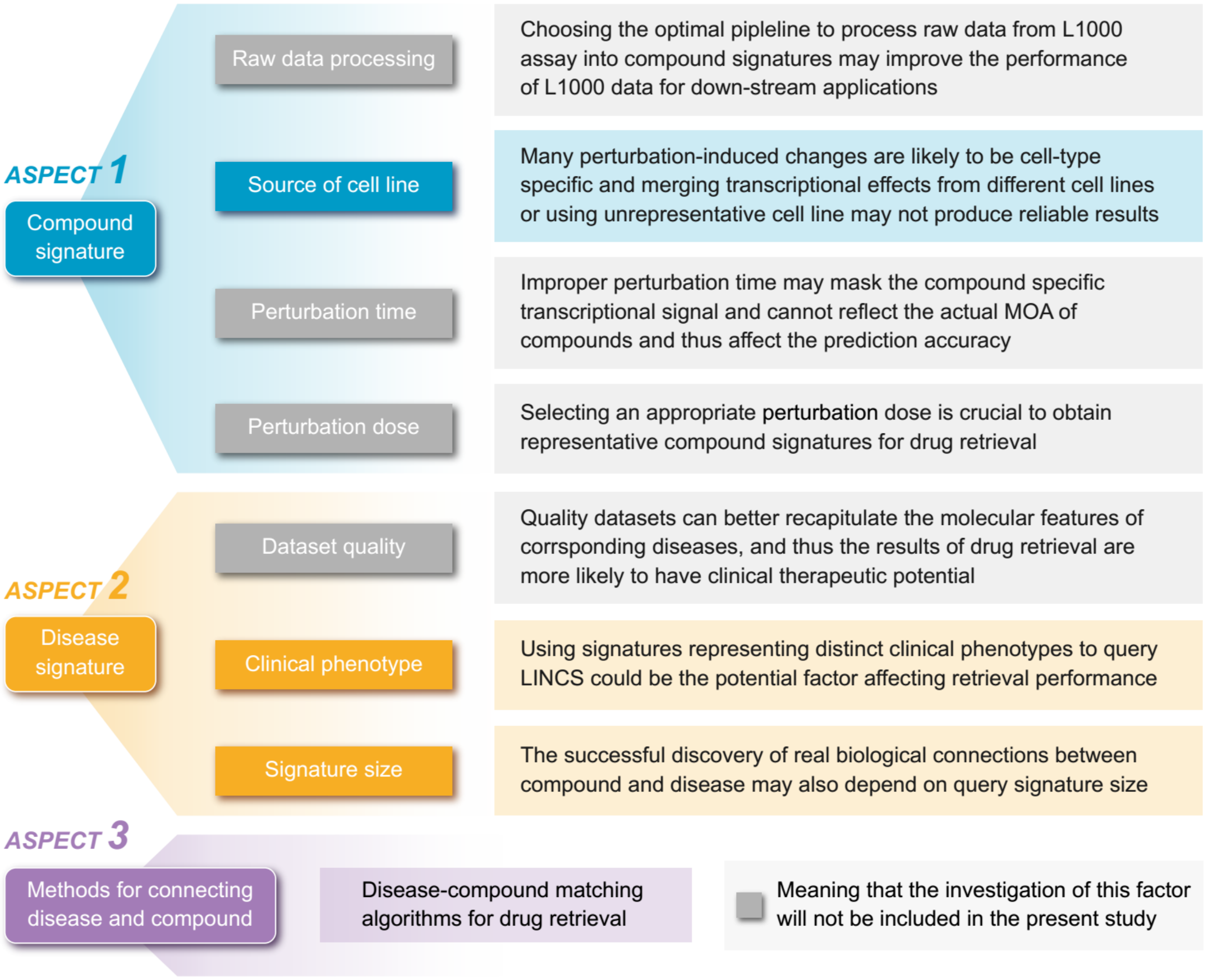


**Fig S1. A summary of potential factors influencing the accuracy of 'signature reversion'-based computational approach.** Within the framework of this approach, there are mainly three components: compound signature, disease signature, and signature matching methods. Each component is likely to be influenced by several factors. In addition to the brief descriptions illustrated in this figure, we also discussed these factors in more detail in Supplementary Discussion.

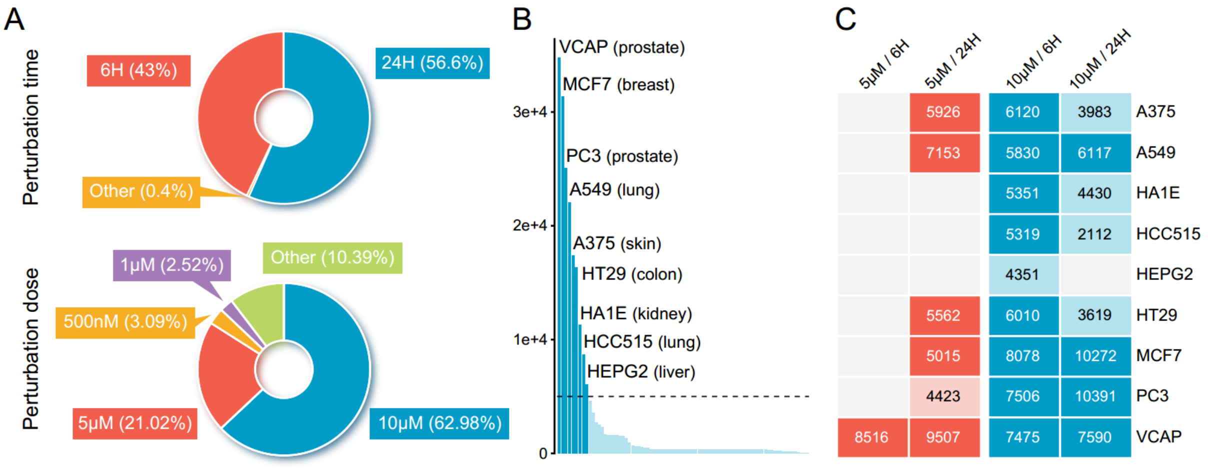


**Fig S2. An overview of compound-induced expression profiles in LINCS. (a)** The distribution of compound profiles of different perturbation time (upper) and concentrations (lower) across all the compound experiments in LINCS dataset. **(b)** The profile count distribution of all 71 cell lines in LINCS. Each bar represents the number of available compound profiles per cell line. The nine most profiled cell lines were labeled in the figure. **(c)** Heat map integrating annotation of the cell lines with perturbation time and concentration. The specific values have not been displayed if there are less than 2,000 profiles in the combination of cell line and experimental conditions.

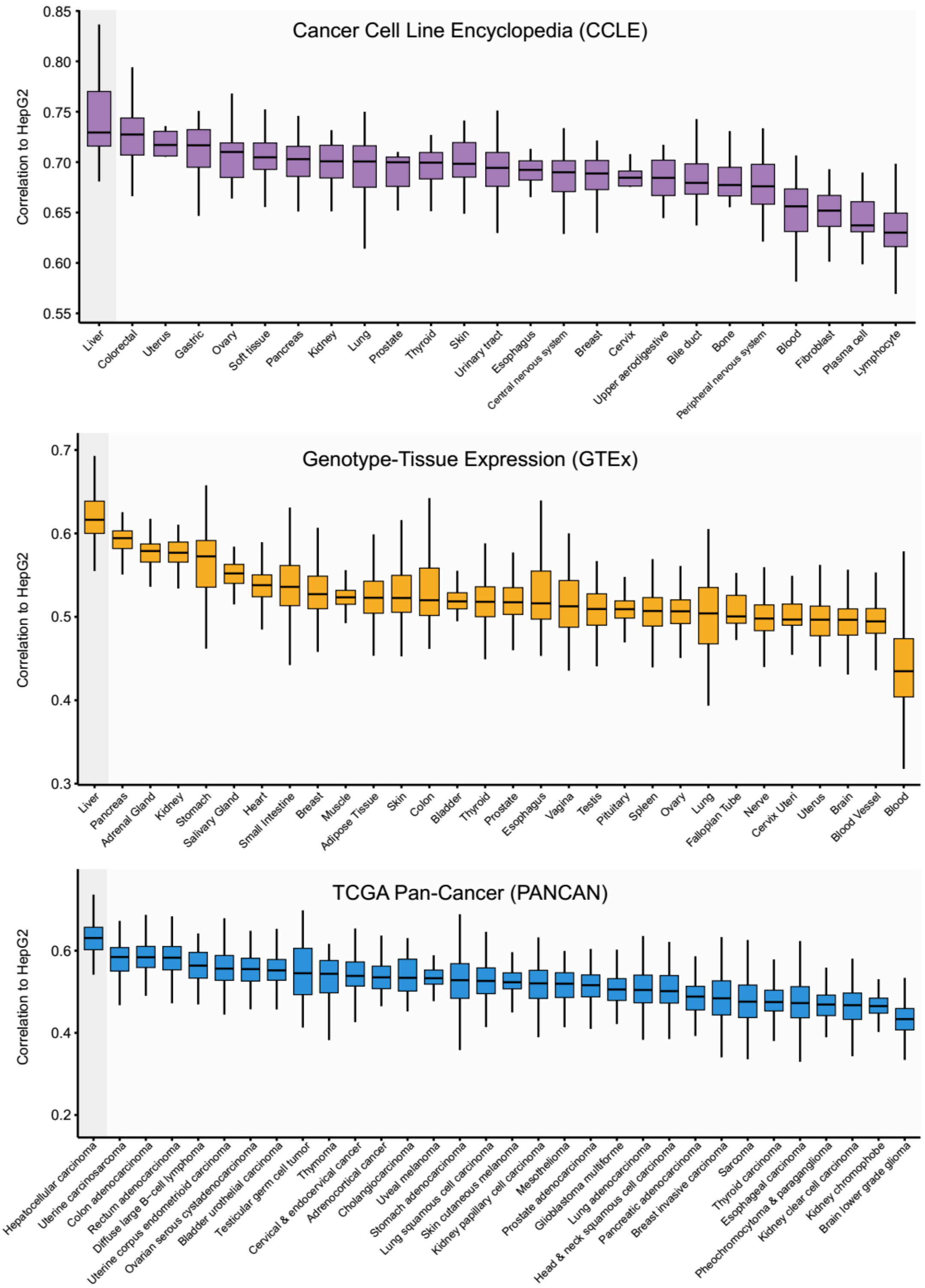


**Fig S3. Correlations between HepG2 cell line and other cancer cell lines or normal/tumor tissues.** Expression data were derived from CCLE, GTEx, and TCGA Pan-Cancer, respectively. Correlations were determined by ranked-based Spearman correlation analysis. The line within the boxes represents the median value, the bottom and top of the boxes denote the interquartile range, and the vertical line represents 1.5 times the interquartile range.

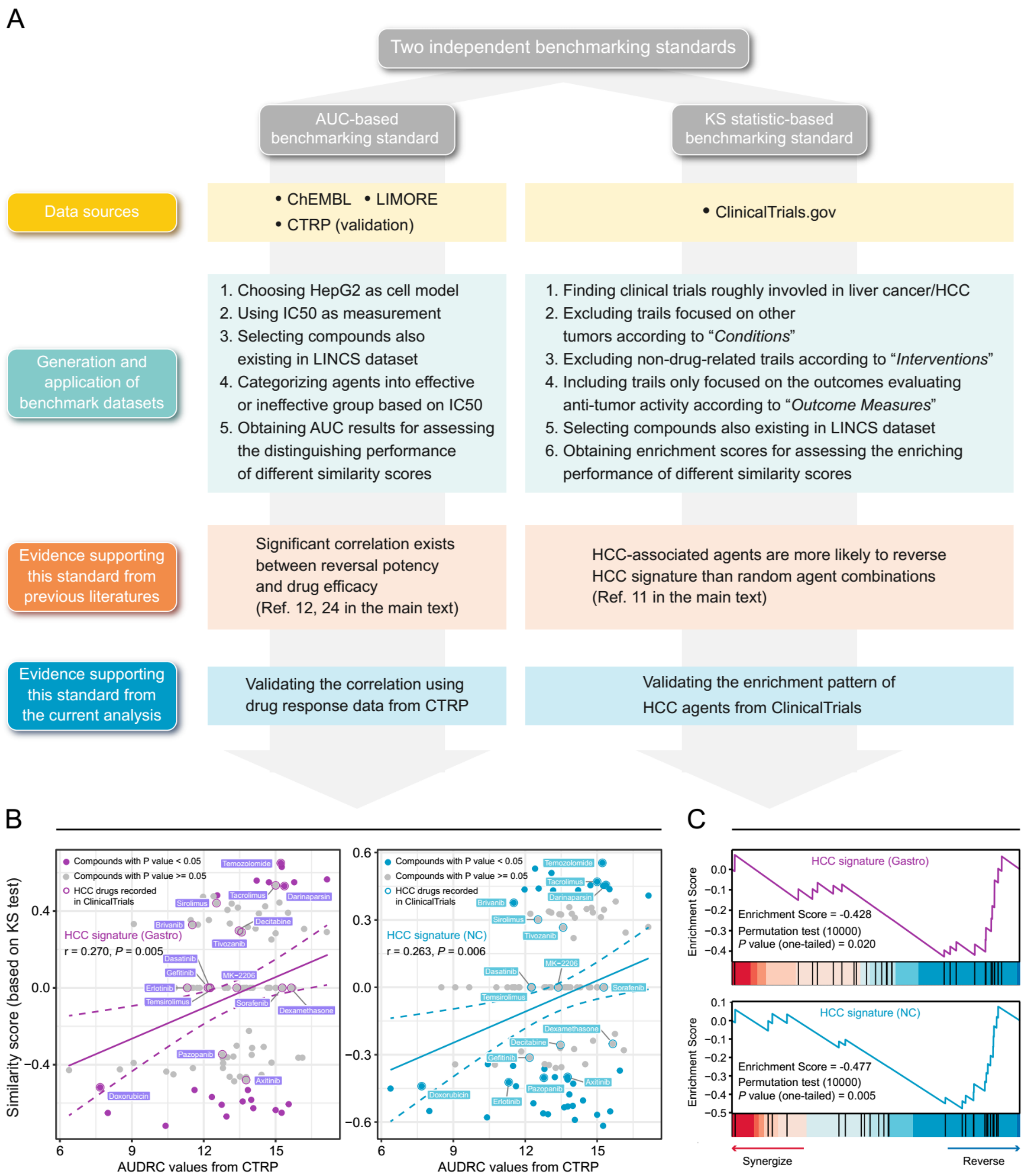


**Fig S4. The procedure for establishing two novel benchmarking standards. (a)** Flow chart of the data collection and hypothesis validation for the AUC-based (left) and KS statistic-based (right) benchmarking standards. **(b)** Correlation between drug efficacy (AUDRC values) and reversal potency (KS-based similarity scores). Two previously published query signatures, including $Sig_{gastro}$ (left) and $Sig_{NC}$ (right), were utilized to calculate similarity scores. Note that lower similarity scores indicate higher reversal potency and lower AUDRC values imply greater drug sensitivity. Color toward gray indicates no statistical significance determined by KS test. **(c)** Reversal potency of HCC agents demonstrated by enrichment analysis. $Sig_{gastro}$ (upper) and $Sig_{NC}$ (lower) were used to compute similarity scores.

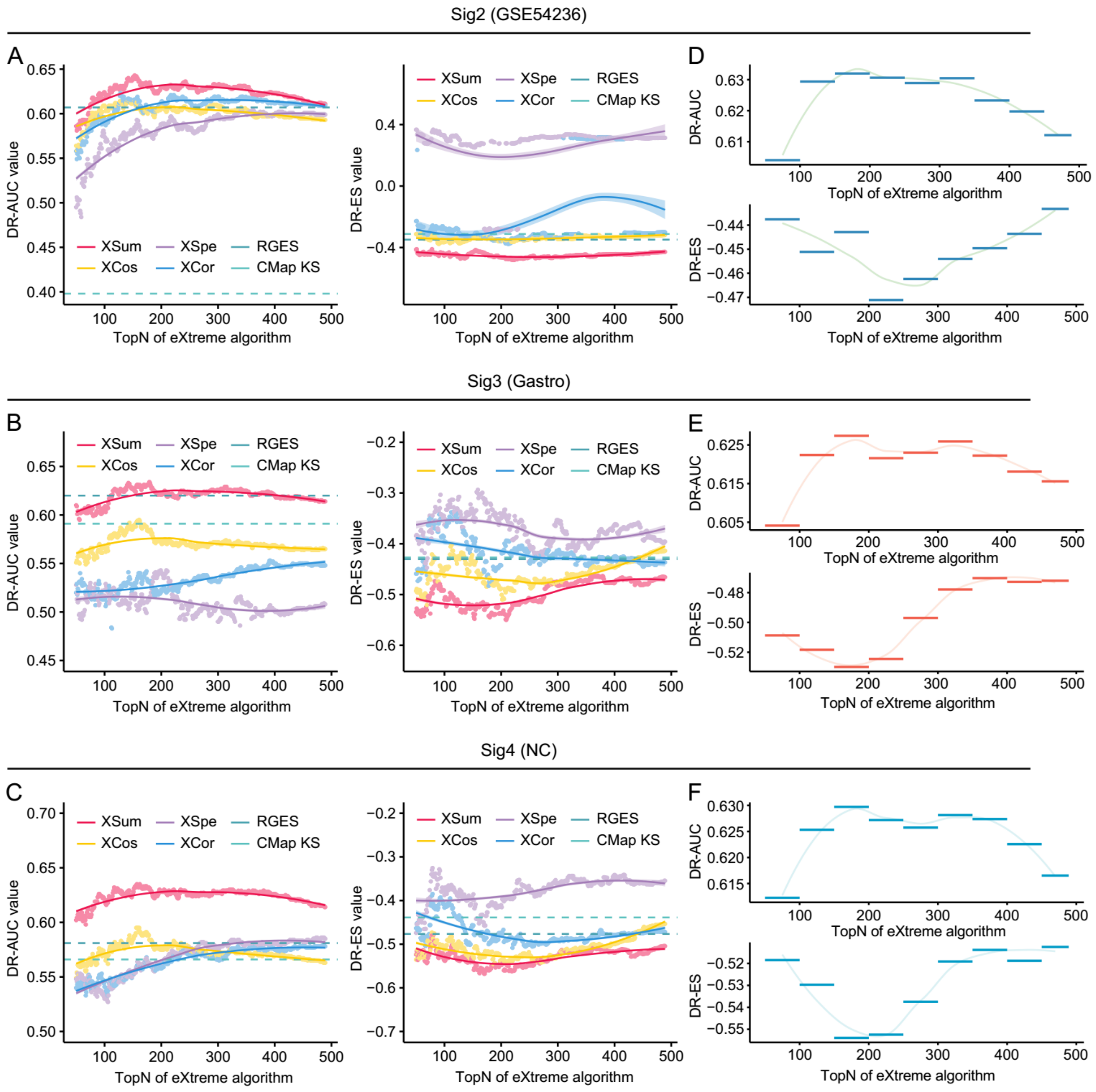


**Fig S5. Benchmarking methodologies and parameters in the conditions of using different query signatures.** Retrieval performance of six matching methods evaluated by AUC-based benchmarking standard (left) and KS statistic-based benchmarking standard (right) in the conditions of using Sig$_{GSE54236}$ **(a)**, Sig$_{gastro}$ **(b)**, and Sig$_{NC}$ **(c)** for querying LINCS. AUC-based (upper) and KS statistic-based (lower) standardized performance measurements of XSum method in the conditions of using Sig$_{GSE54236}$ **(d)**, Sig$_{gastro}$ **(e)**, and Sig$_{NC}$ **(f)** as query signatures.

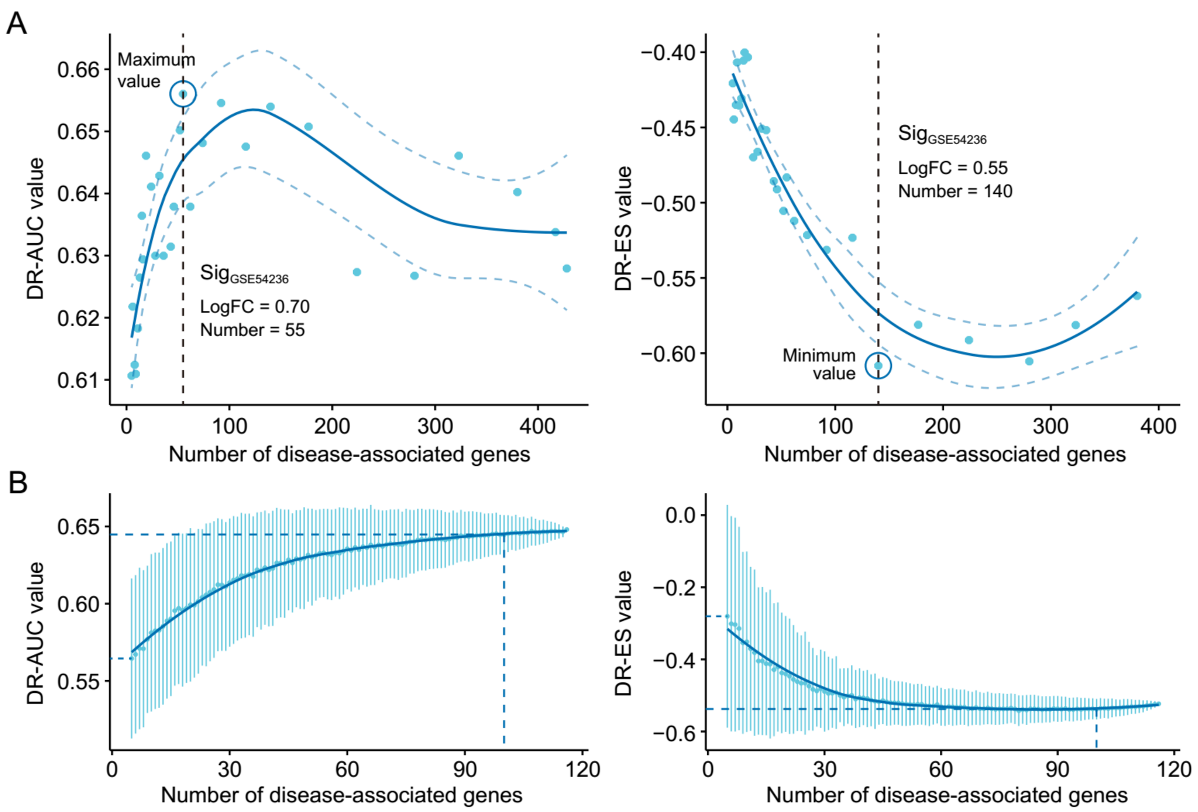


**Fig S6. The influences of query signature size on retrieval performance.** Relationship between query signature size determined by iterative fold change-based approach **(a)** or random sampling-based approach **(b)** and retrieval performance evaluated by AUC-based standard (left) and KS statistic-based standard (right). GSE54236 cohort was used for generating candidate query signatures for evaluation.

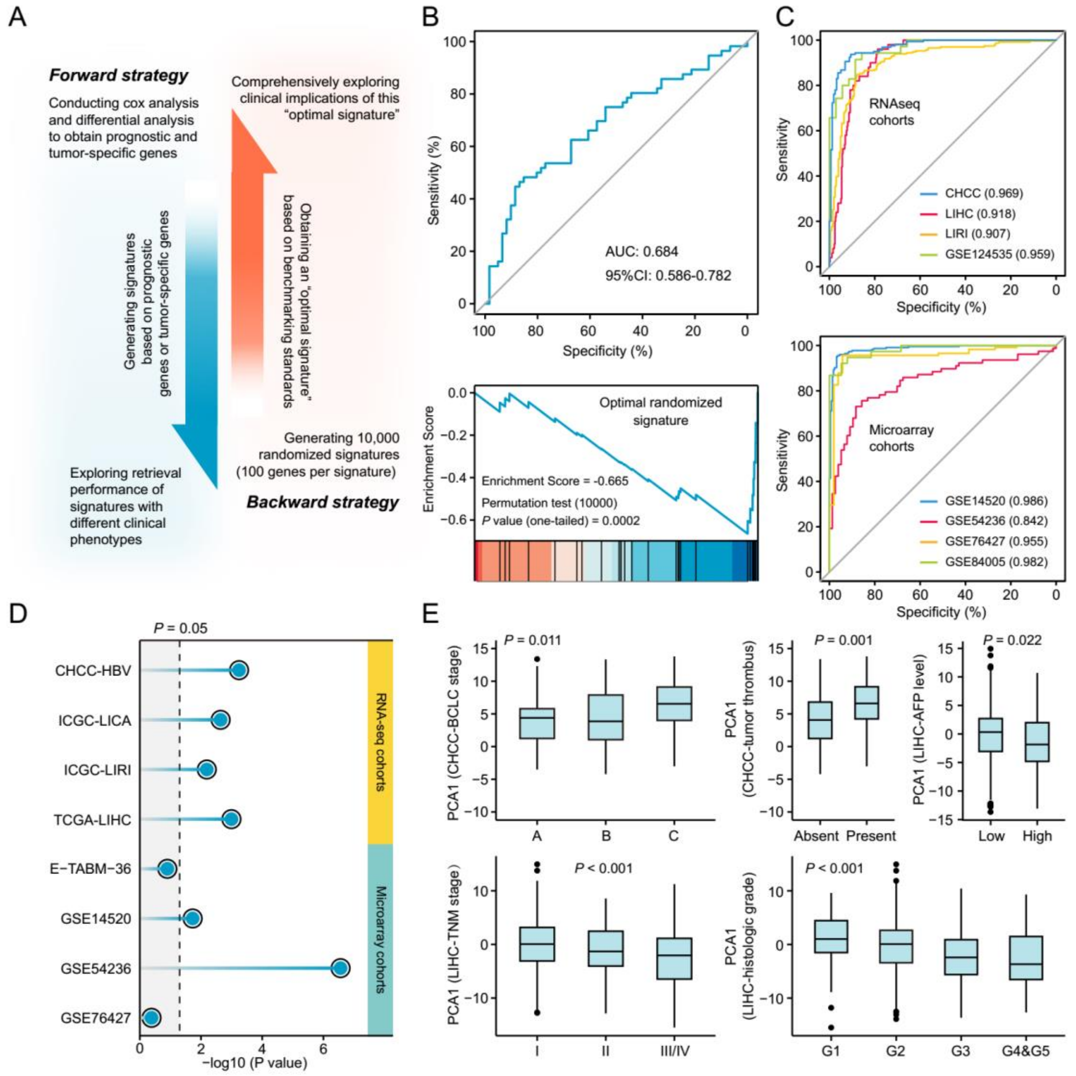


**Fig S7. Investigating the necessary properties that quality query signatures should possess. (a)** Schematic illustration of forward and backward strategy adopted to investigate whether the factor associated with clinical phenotype of query signature can affect computational therapeutic discovery. **(b)** The DR-AUC value and DR-ES value of the optimal randomized signature showed by ROC curve (upper) and enrichment plot (lower). **(c)** The association between the optimal signature and the clinical phenotype of discordant expression pattern suggested by ROC curves based on RNA sequencing cohorts (upper) and Microarray cohorts (lower). **(d)** The association between the optimal signature and the clinical phenotype of prognosis. Color toward gray indicates no statistical significance. **(e)** The association between the optimal signature and multiple clinical characteristics, including BCLC stage, tumor thrombus, AFP level, TNM stage, and histologic grade. The statistical significance of difference between groups was determined using either Kruskal-Wallis or Wilcoxon sum rank tests.

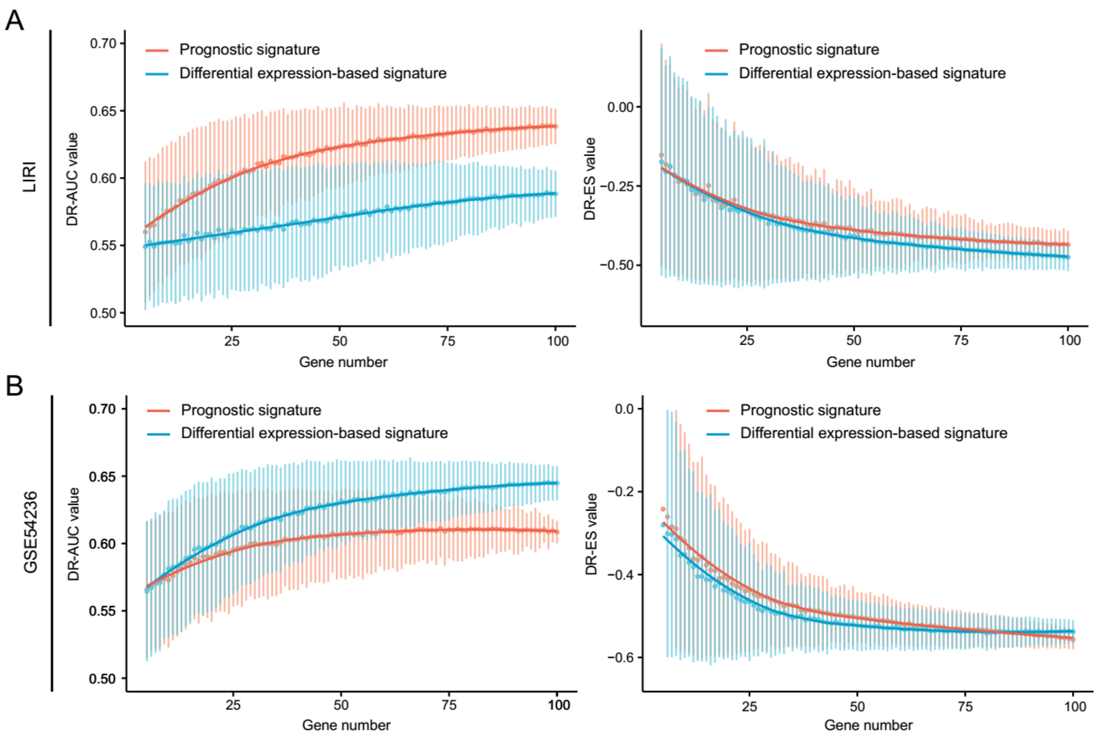


**Fig S8. The influences of query signature phenotype on retrieval performance.** Comparison of retrieval performance evaluated by AUC-based standard (left) and KS statistic-based standard (right) between prognosis-associated signatures and discordant expression-associated signatures generated by LIRI cohort **(a)** or GSE54236 cohort **(b)**.

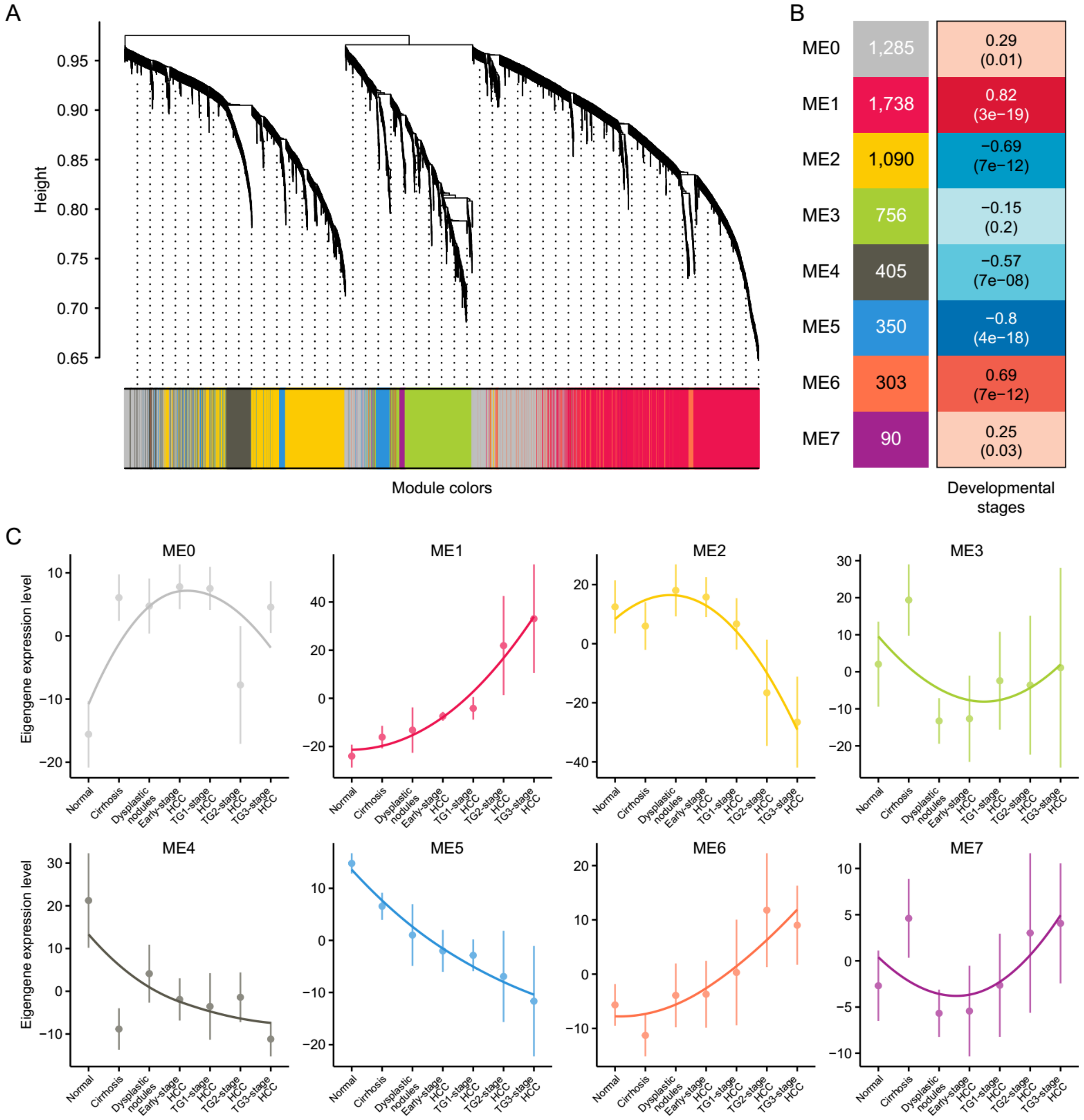


**Fig S9. Weighted gene co-expression network analysis. (a)** Hierarchical cluster tree showing eight modules of co-expressed genes identified by WGCNA. Each of the stage-associated genes is represented by a leaf in the tree, and each of the eight modules by a major tree branch. **(b)** Module-trait (developmental stage) relationship and corresponding *P* values. The left panel shows the eight modules and the number of genes in each module, and the colour scale on right shows module-trait correlation from –1 (blue) to 1 (red). **(c)** The expression pattern of eight modules. Note that grey module represents unassigned genes.

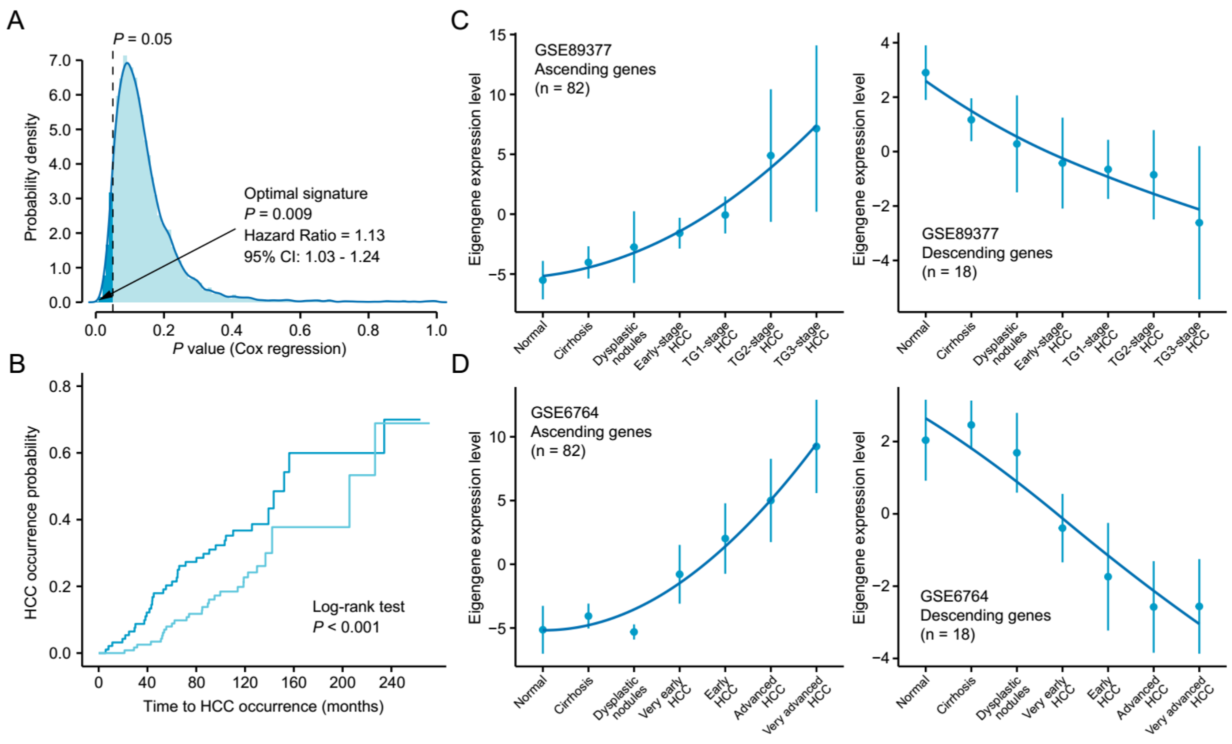


**Fig S10. Identification and validation of the novel query signature. (a)** The distribution of the statistical significance results of 10,000 cox proportional hazards regression analysis. The optimal signature with the most significant relevance to HCC occurrence was labeled in the figure. **(b)** Kaplan-Meier cumulative hazard rates for HCC occurrence according to the groups determined by the optimal signature expression. **(c)** Validation of the expression pattern of the 'ascending' (left) and 'descending' (right) module based on the optimal signature in GSE89377 cohort. **(d)** Validation of the expression pattern of the 'ascending' (left) and 'descending' (right) module based on the optimal signature in GSE6764 cohort.

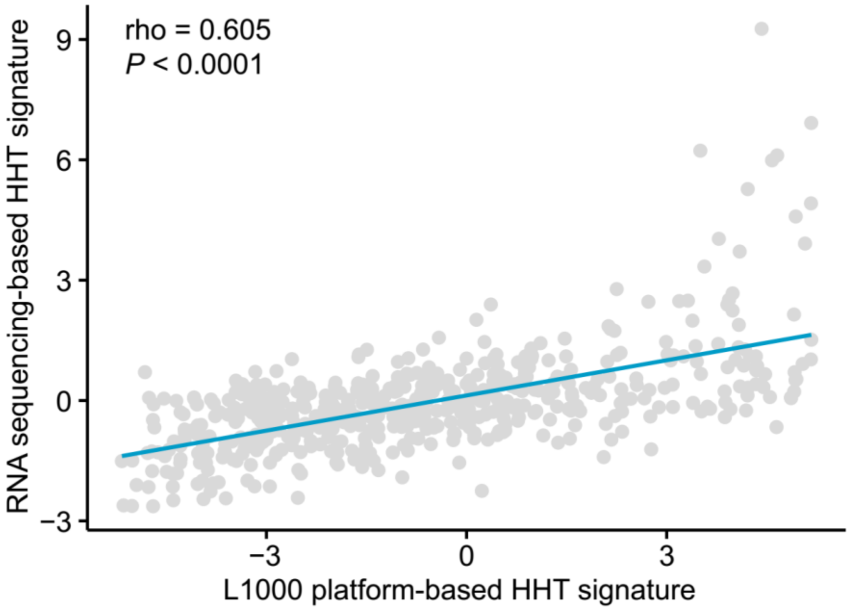


**Fig S11. Assessment of the reliability of L1000 platform based HHT signature.** We treated HepG2 cells with the same conditions in LINCS (perturbation time: 6 hours, perturbation dose: 10μM) and utilized RNA sequencing to profile the HHT-induced expression changes. Association between RNA sequencing-based HHT signature and L1000 platform-based HHT signature were then determined by ranked-based Spearman correlation.

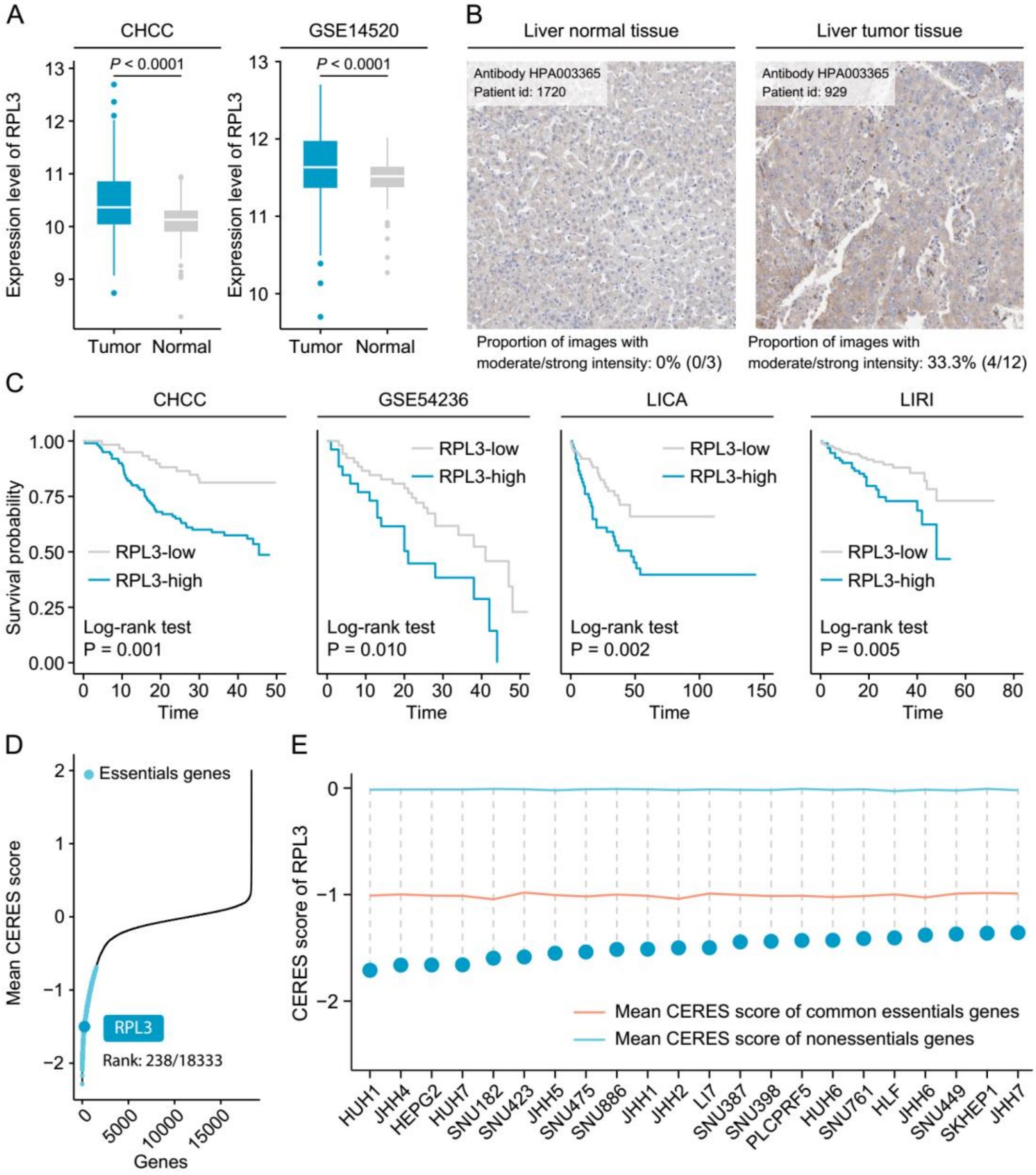


**Fig S12. Clinical and biological characterization of RPL3 in liver cancer. (a)** The comparison of mRNA expression level of RPL3 between tumor and normal tissues in CHCC (left) and GSE14520 (right) cohorts. Statistical significance of difference was determined using Wilcoxon rank-sum test. **(b)** Representative images of immunohistochemical staining of RPL3 in liver normal (left) and tumor tissues (right) from the Human Protein Atlas (HPA) program. **(c)** Comparison of survival curves between high RPL3 expression and low RPL3 expression groups. **(d)** Distribution of gene dependency score (CERES score) of 18,333 protein coding genes in liver cancer cell lines. A lower CERES score of certain gene indicates a higher likelihood that this gene is essential in cell growth and survival. **(e)** The gene dependency of RPL3 across 22 liver cancer cell lines.

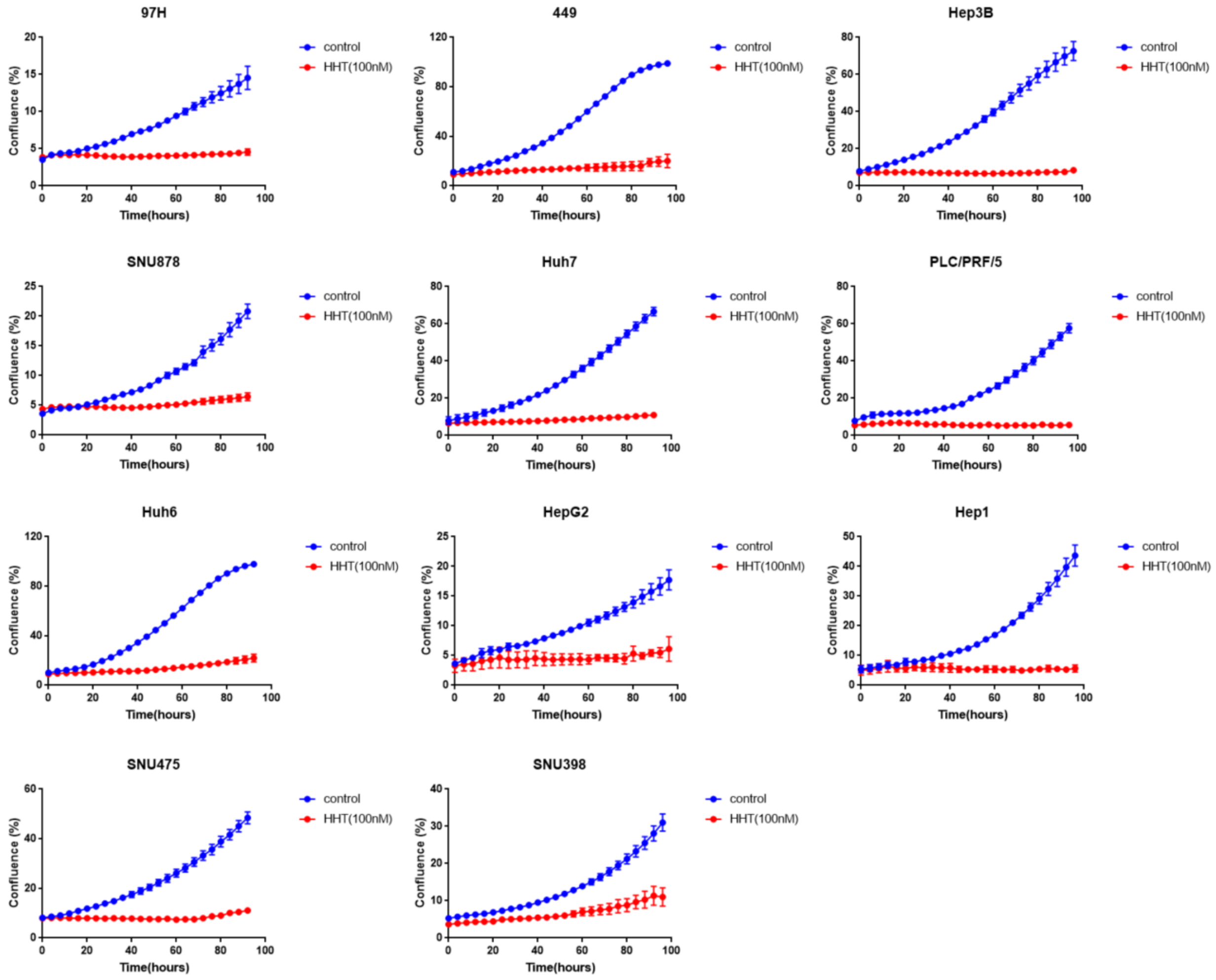


**Fig S13. The effect of HHT on cell proliferation across 11 liver cancer cell lines.** Cell proliferation rate was analyzed by IncuCyte ZOOM system every four hours for 100 hours.

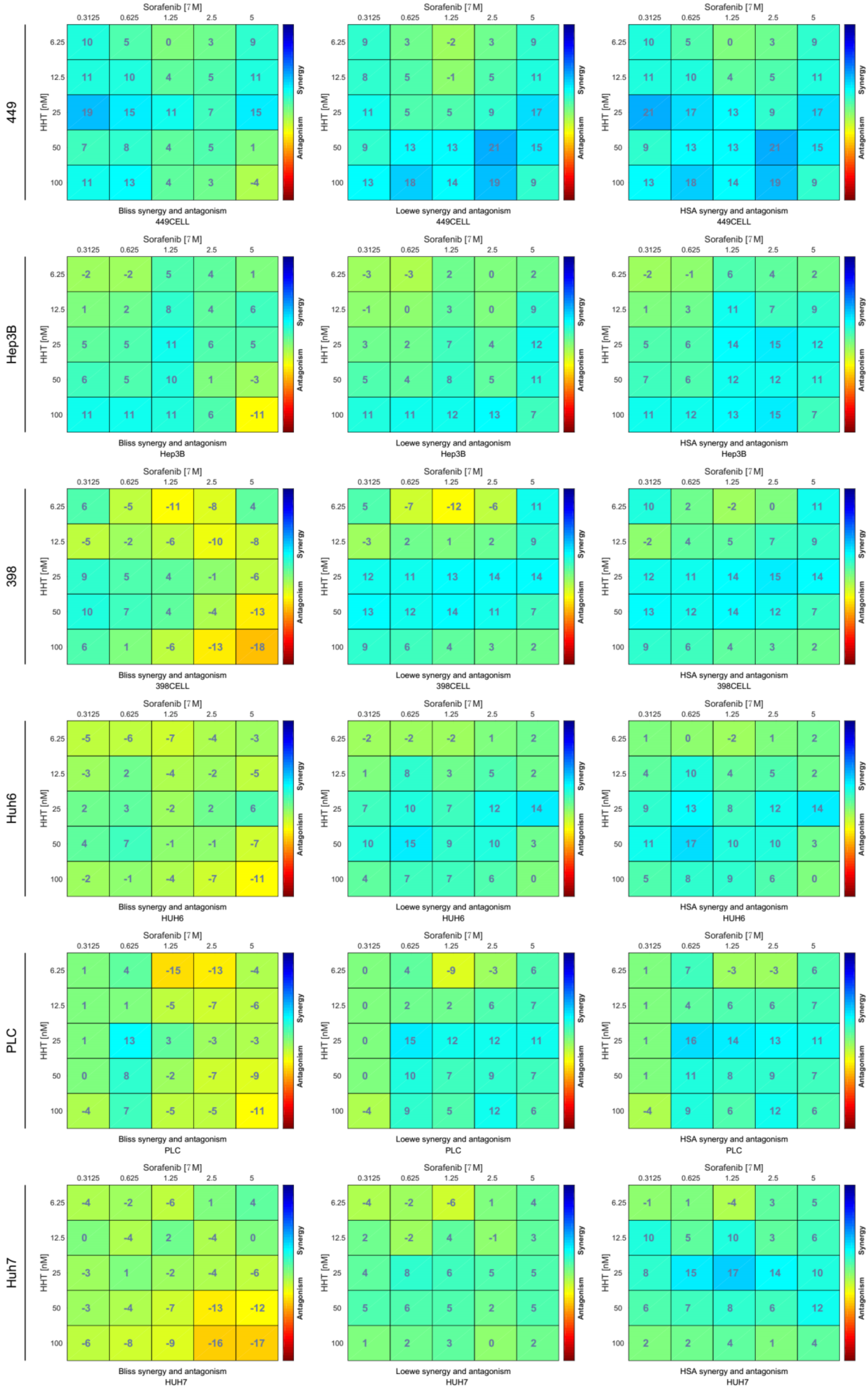
449
Sorafenib [7M]
0.3125 0.625 1.25 2.5 5
HHT [nM]
6.25 10 5 0 3 9
12.5 11 10 4 5 11
25 19 15 11 7 15
50 7 8 4 5 1
100 11 13 4 3 -4
Synergy
Antagonism
Bliss synergy and antagonism
449CELL
Sorafenib [7M]
6.25 9 3 -2 3 9
12.5 8 5 -1 5 11
25 11 5 5 9 17
50 9 13 13 21 15
100 13 18 14 19 9
Loewe synergy and antagonism
449CELL
Sorafenib [7M]
6.25 10 5 0 3 9
12.5 11 10 4 5 11
25 21 17 13 9 17
50 9 13 13 21 15
100 13 18 14 19 9
HSA synergy and antagonism
449CELL
Hep3B
6.25 -2 -2 5 4 1
12.5 1 2 8 4 6
25 5 5 11 6 5
50 6 5 10 1 -3
100 11 11 11 6 -11
Bliss synergy and antagonism
Hep3B
6.25 -3 -3 2 0 2
12.5 -1 0 3 0 9
25 3 2 7 4 12
50 5 4 8 5 11
100 11 11 12 13 7
Loewe synergy and antagonism
Hep3B
6.25 -2 -1 6 4 2
12.5 1 3 11 7 9
25 5 6 14 15 12
50 7 6 12 12 11
100 11 12 13 15 7
HSA synergy and antagonism
Hep3B
398
6.25 6 -5 -11 -8 4
12.5 -5 -2 -6 -10 -8
25 9 5 4 -1 -6
50 10 7 4 -4 -13
100 6 1 -6 -13 -18
Bliss synergy and antagonism
398CELL
6.25 5 -7 -12 -6 11
12.5 -3 2 1 2 9
25 12 11 13 14 14
50 13 12 14 11 7
100 9 6 4 3 2
Loewe synergy and antagonism
398CELL
6.25 10 2 -2 0 11
12.5 -2 4 5 7 9
25 12 11 14 15 14
50 13 12 14 12 7
100 9 6 4 3 2
HSA synergy and antagonism
398CELL
Huh6
6.25 -5 -6 -7 -4 -3
12.5 -3 2 -4 -2 -5
25 2 3 -2 2 6
50 4 7 -1 -1 -7
100 -2 -1 -4 -7 -11
Bliss synergy and antagonism
HUH6
6.25 -2 -2 -2 1 2
12.5 1 8 3 5 2
25 7 10 7 12 14
50 10 15 9 10 3
100 4 7 7 6 0
Loewe synergy and antagonism
HUH6
6.25 1 0 -2 1 2
12.5 4 10 4 5 2
25 9 13 8 12 14
50 11 17 10 10 3
100 5 8 9 6 0
HSA synergy and antagonism
HUH6
PLC
6.25 1 4 -15 -13 -4
12.5 1 1 -5 -7 -6
25 1 13 3 -3 -3
50 0 8 -2 -7 -9
100 -4 7 -5 -5 -11
Bliss synergy and antagonism
PLC
6.25 0 4 -9 -3 6
12.5 0 2 2 6 7
25 0 15 12 12 11
50 0 10 7 9 7
100 -4 9 5 12 6
Loewe synergy and antagonism
PLC
6.25 1 7 -3 -3 6
12.5 1 4 6 6 7
25 1 16 14 13 11
50 1 11 8 9 7
100 -4 9 6 12 6
HSA synergy and antagonism
PLC
Huh7
6.25 -4 -2 -6 1 4
12.5 0 -4 2 -4 0
25 -3 1 -2 -4 -6
50 -3 -4 -7 -13 -12
100 -6 -8 -9 -16 -17
Bliss synergy and antagonism
HUH7
6.25 -4 -2 -6 1 4
12.5 2 -2 4 -1 3
25 4 8 6 5 5
50 5 6 5 2 5
100 1 2 3 0 2
Loewe synergy and antagonism
HUH7
6.25 -1 1 -4 3 5
12.5 10 5 10 3 6
25 8 15 17 14 10
50 6 7 8 6 12
100 2 2 4 1 4
HSA synergy and antagonism
HUH7

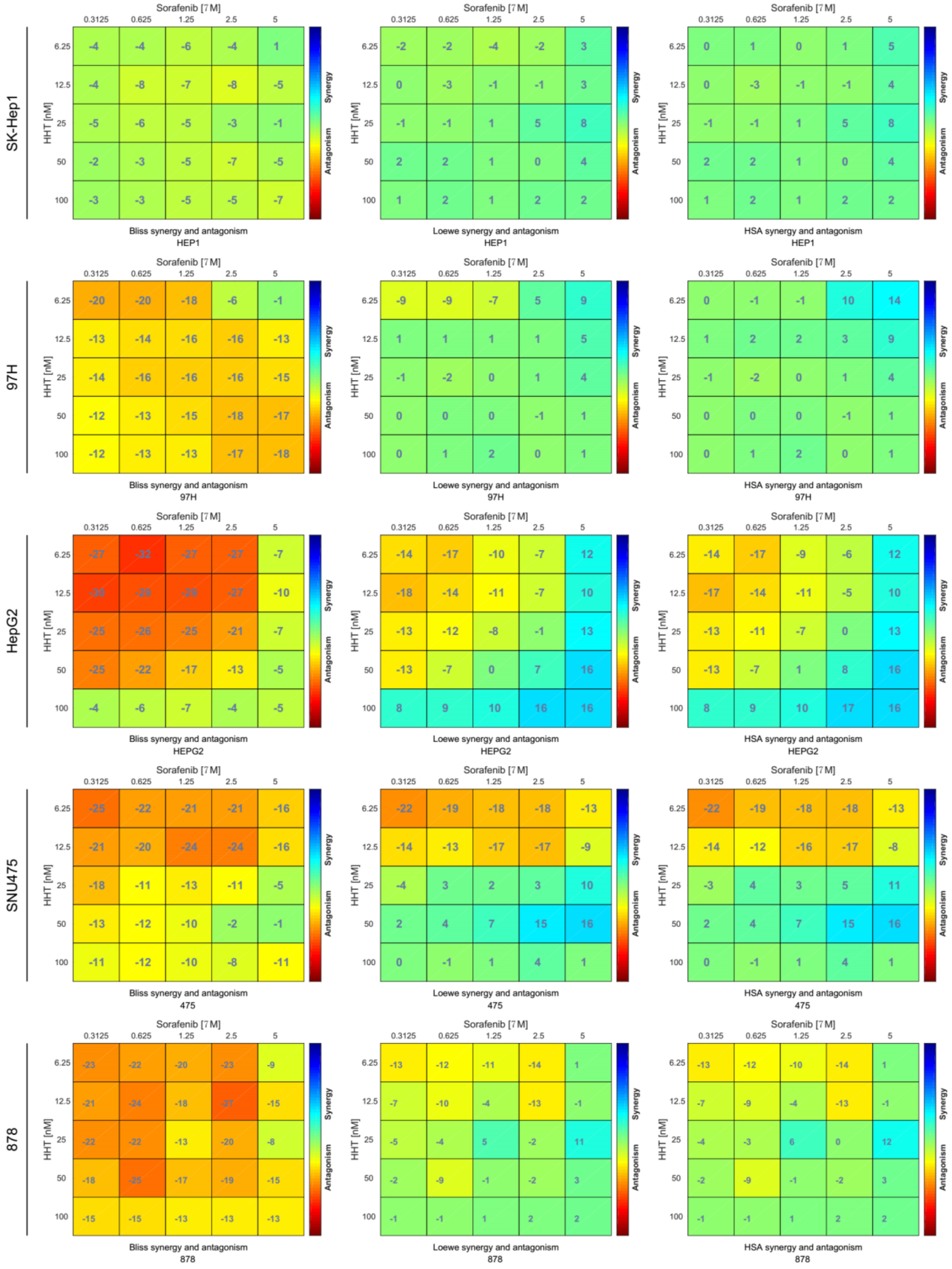

**Fig S14. The anti-tumor effect of the combination of HHT and sorafenib across 11 different liver cancer cell lines.** Three different models, including Bliss, Loewe, and HAS models, were adopted for measuring synergistic effect. A higher positive score indicated stronger drug synergy.

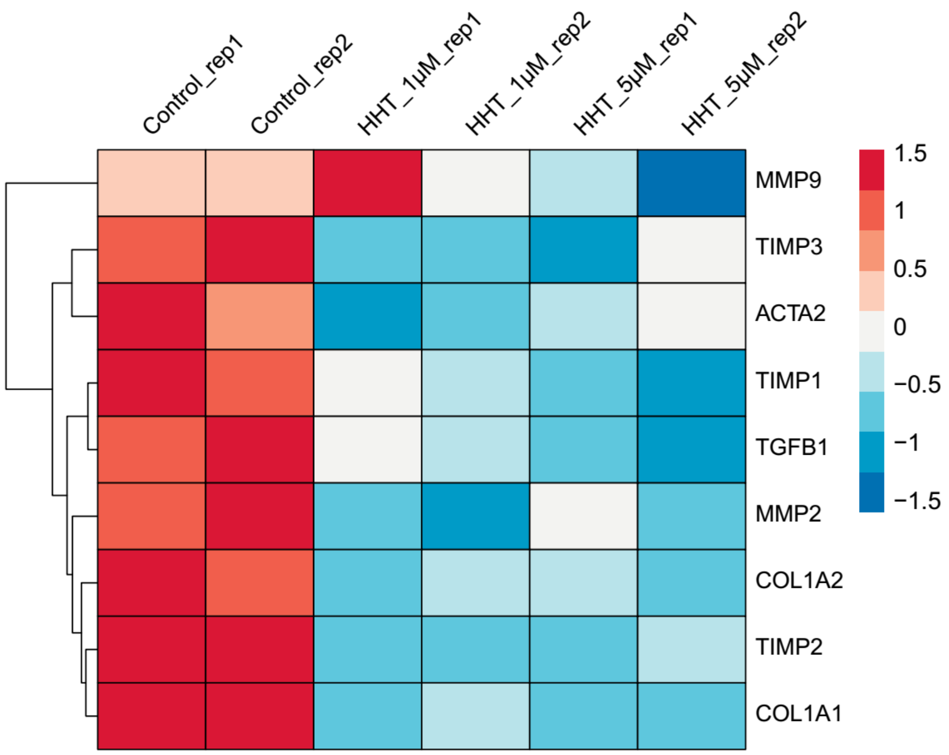


**Fig S15.** Comparison of the expression of nine fibrosis-associated genes between control LX2 and HHT-treated LX2.
